## Supporting information for "Contact networks have small metric backbones that maintain community structure and are primary transmission subgraphs"

### S1 Text. Additional method details and results on other contact networks.

#### A Network Construction

In the main manuscript we focused our results on a *social* aggregation and normalization of contact data into a contact relation graph $R(X)$ used in Section 2. The social normalization is a per-individual accounting of the number of interactions each individual had during the measurement. This aggregation of social contact data is intuitive and well suited to social contexts, as it weighs individuals separately, possibly accounting for inter-individual social variability. However, different normalizations are possible, ultimately resulting in different proximity and distance graphs. We considered two additional normalizations to ascertain that the results are robust to aggregation procedure.

For each dataset, contact events between individuals $x_i$ and $x_j$ from population $X$ are represented in a symmetric graph $R_w(X)$, where $w$ denotes a chosen time window, with $i, j, w \in \mathbb{N}$. While one can explore larger time windows, we used the smallest time window resolution available for each dataset, typically $w = 20$ seconds. Thus, henceforth and in manuscript we refer to the contact graph as simply $R(X)$. Entries of its adjacency matrix $r_{ij}$ denote the number of time windows in which individuals $x_i$ and $x_j$ were observed in close proximity. The distinct normalizations depend only on the diagonal entries of the adjacency matrix, $r_{ii}$. In the case of *social* normalization we have $r_{ii} = \sum_{j \neq i} r_{ij}$. That is, $r_{ii}$ denotes the total number of time windows in which individual $x_i$ was observed in a close social interaction with any other individual $x_j \in X$. In the *individual time* normalization, $r_{ii}$ denotes the total number of time windows in which individual $x_i$ was observed in the experiment, thus accounting for the potential of an individual influence to spread by the amount of time he or she is present in the experiment. Lastly, in the *experiment time* normalization $r_{ii}$ is, for individuals $x_i$, the total number of time windows in the entire experiment.

From the different contact graphs for each normalization, proximity, $P(X)$, and distance, $D(X)$, graphs are obtained via Eqs 1 and 2 in Section 2, respectively. Fig A shows the distribution of distance and proximity weights for two SocioPatterns datasets: primary school (Fr-PS) [45] and hospital (Fr-Ho) [42]. Both the *social* and *individual time* normalization are alike (left and middle plots, respectively), with differences at the extremes. Conversely, the *experiment time* normalization (right plots) differs from the others. However, as shown in the tables for each dataset in Section C, the size of the metric backbone changes very little between the various normalizations. Therefore, in the manuscript we present results for the social normalization.

The code used to compute networks from social contact data is available at https://github.com/rionbr/SocialBackbone.

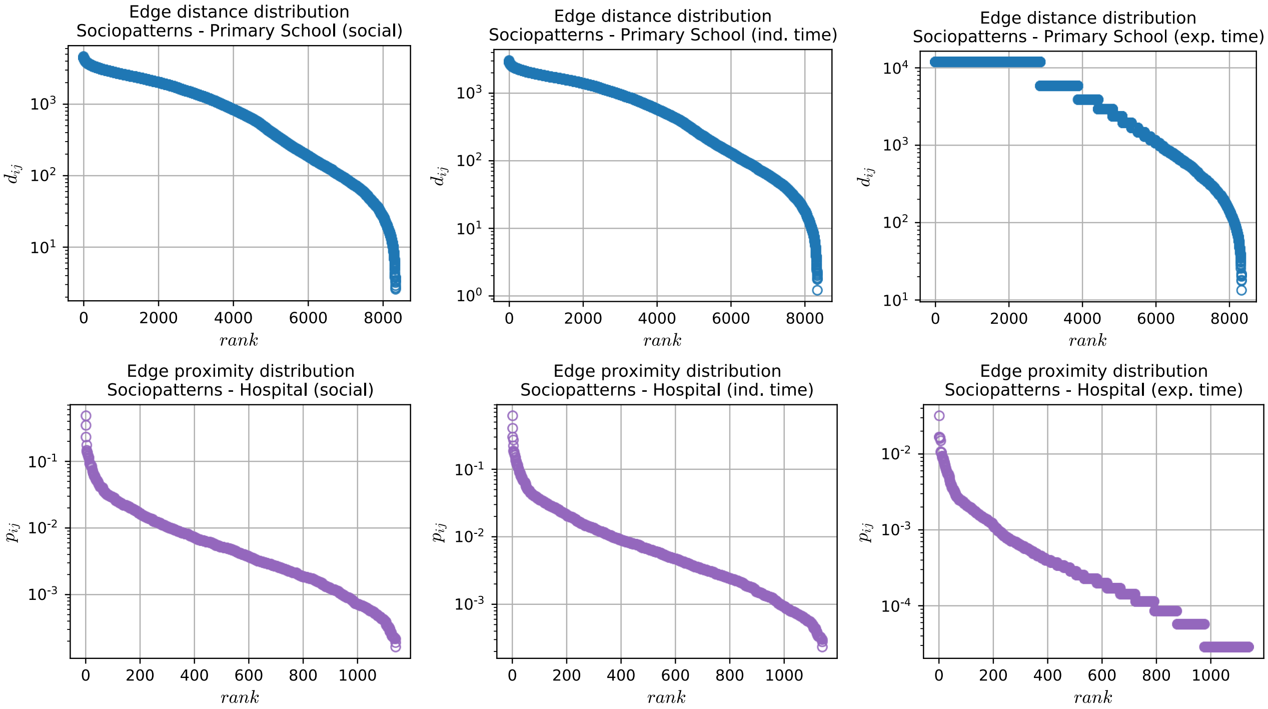


**Fig A. Edge distributions for *social* (left), *individual time* (middle), and *experiment time* (right) normalizations.** Distance, $d_{ij}$, and proximity, $p_{ij}$, distributions are shown for SocioPatterns datasets *Primary school* (Fr-PS) (top) and *Hospital* (Fr-Ho) (bottom), respectively. For visual representation distances $d_{ij}$ are shown log converted.

### B Synthetic networks

To better understand how well the metric backbone captures community structure in a social network, we use stochastic block models to control how much class metalabels match the underlying community structure—something we cannot control in the real-world contact networks as discussed in main manuscript (Section 3.1). Here we detail the construction and analysis of two ensembles of synthetic networks built from stochastic block models. We start by synthesizing networks with 160 nodes and 4 modules (A, B, C, and D), each containing 40 nodes each (see Fig BA). We can think that the nodes may represent students in classrooms, though they can represent other social associations. The underlying connectivity of the networks is defined by a predefined Stochastic Block Model (SBM) [51]. SBMs were first proposed for social network analysis in the early 80s and are still extensively used to generate random networks with modular group tendencies [1]. The parameters of the SBM are the size of each module, and a module probability matrix that defines the likelihood that nodes in a module connect to nodes in the same module and in the other modules (see Fig BA & D1 for an example).

We consider two distinct SBM to study the cases of more or less defined community structure. Thus, one SBM is defined by '*high*' connectivity (Fig BD1), and the other by '*low*' connectivity (Fig BC1) Also note that classes C and D have a hierarchical connectivity structure, given by the predefined SBM. Here the interpretation is that students in these classrooms belong to the same grade and thus share a number of classes and interests. The proximity $p_{ij}$ weights for each edge between $x_i$ and $x_j$—and thus also the isomorphic distance $d_{ij}$ via map $\varphi$ (Section 2)—are sampled from a real contact network, the SocioPatterns French Primary School (Fr-PS; details in Section C.1). Importantly, edges are sampled taking into consideration if they connect two students belonging to the *same classroom* or *across different classrooms* in the Fr-PS. This provide us a realistic differentiation of *within module* and *across module* connectivity weights (see Fig BB). An instance of the adjacency matrix of a generated network and its metric backbone can be seen in Fig BC2 & D2, for the '*low*' and '*high*' connectivity models, respectively. Their network visualizations, plotted using a force layout algorithm, can also be seen in Fig BC3 & D3 and Fig BC4 & D4. Finally, we ran Louvain and Infomap to estimate network modules and computed the same module similarity measures described in the main manuscript (see Section 4.2). In Table A we present network statistics for both models, while in Tables B and C we show the similarity measure results. To estimate variability, all results are based on 10 realizations of each SBM model. To compute results for the random backbone, 100 iterations (each with 10 network

**Table A. Synthetic network metric backbone statistics.** Results based on 10 realizations of each network network ('*low*' and '*high*' connectivity). See also Fig B.

| | '*low*' connectivity | '*high*' connectivity |
|---|---|---|
| $D(X)$ | | |
| Nodes | 160 | 160 |
| Edges | 1,560±36 | 3,520±34 |
| $B(X)$ | | |
| Metric edges ($s_{ij} = 1$) | 409±19 (26±1%) | 503±14 (14±0.4%) |
| Semi-metric edges ($s_{ij} > 1$) | 1,152±36 (74±1%) | 3,017±37 (86±0.4%) |

realizations) were used, as performed for the real datasets in the main manuscript.

The synthetic networks generated from both SBM allows us to draw some important conclusions. First, for both low and high connectivity, the metric backbone preserves well the community structure of the original network. This can be observed by inspecting the adjacency matrix and the network visualization (see Fig BC2-4 and Fig BD2-4). Naturally, the high connectivity case results in more defined community structure for networks and their subgraphs. But the community structure detected on the metric backbone subgraphs is systematically more similar to the one detected on the full generated networks, than the community structures of threshold and random subgraphs of the same size. This is so for both the high and low connectivity SBM, for Louvain and Infomap community structure detection, and across most measures of modularity similarity (see Tables B and C).

It is also noteworthy that Louvain is more accurate than Infomap in detecting the SBM modules in the generated networks and their subgraphs for both the low and the high connectivity models. For instance, for the '*low*' connectivity model, while Louvain returns 4.3±0.5 and 6.1±1.6 modules respectively for the original networks and their metric backbone, Infomap finds 9.1±2 and 19±3.6 modules. Even in the naturally easier '*high*' connectivity model, the 4 SBM modules are correctly found by Louvain for every generated original network and their metric backbone, while Infomap returns 6.7±1.42 modules for the metric backbone (see Table B). Thus, in the main manuscript we show only results for Louvain, since Infomap is prone to detect many more modules than the underlying community structure in the SBMs.

**Table B. '*High*' connectivity synthetic network measures of module similarity.** Values for 10 randomly generated network realizations; random subgraphs based on 100 instances. See also Fig B.

| | | Original | Metric | Threshold | Random |
|---|---|---|---|---|---|
| | $m$ | | | | |
| | Metalabels | 4 | - | - | - |
| | Louvain | 4 | 4 | 4.1±0.32 | 11±1.5 |
| | Infomap | 4 | 6.7±1.42 | 7.4±1.78 | 29±3.2 |
| Louvain | $y_{AB}$ | | | | |
| | Metalabels | 1.00±0.00 | 0.99±0.01 | 0.98±0.03 | 0.58±0.04 |
| | Original | - | 0.99±0.01 | 0.98±0.03 | 0.58±0.04 |
| | $J_{A\to B}/J_{B\to A}$ | | | | |
| | Metalabels | 1.00±0.00/1.00±0.00 | 0.99±0.01/0.99±0.01 | 0.99±0.01/0.97±0.06 | 0.57±0.08/0.34±0.05 |
| | Original | - | 0.99±0.08/0.99±0.01 | 0.99±0.01/0.97±0.06 | 0.57±0.08/0.34±0.05 |
| | $h_{A\to B}/h_{B\to A}$ | | | | |
| | Metalabels | 0.00±0.00/0.00±0.00 | 0.02±0.02/0.02±0.02 | 0.02±0.02/0.02±0.02 | 0.40±0.05/0.11±0.05 |
| | Original | - | 0.02±0.02/0.02±0.02 | 0.02±0.02/0.02±0.03 | 0.40±0.05/0.11±0.05 |
| | *CluSim* | | | | |
| | Metalabels | 1.00±0.00 | 0.98±0.01 | 0.99±0.01 | 0.47±0.08 |
| | Original | - | 0.98±0.01 | 0.99±0.01 | 0.47±0.08 |
| | *Adjusted Rand Index* | | | | |
| | Metalabels | 1.00±0.00 | 0.98±0.02 | 0.98±0.02 | 0.xx±0.02 |
| | Original | - | 0.98±0.02 | 0.98±0.02 | 0.xx±0.02 |
| Infomap | $y_{AB}$ | | | | |
| | Metalabels | 1.00±0.00 | 0.78±0.08 | 0.74±0.09 | 0.37±0.02 |
| | Original | - | 0.78±0.08 | 0.74±0.09 | 0.37±0.02 |
| | $J_{A\to B}/J_{B\to A}$ | | | | |
| | Metalabels | 1.00±0.00/1.00±0.00 | 0.90±0.05/0.62±0.13 | 0.88±0.06/0.56±0.13 | 0.30±0.06/0.13±0.02 |
| | Original | - | 0.90±0.05/0.62±0.13 | 0.88±0.06/0.56±0.13 | 0.30±0.06/0.13±0.02 |
| | $h_{A\to B}/h_{B\to A}$ | | | | |
| | Metalabels | 0.00±0.00/0.00±0.00 | 0.13±0.05/0.01±0.02 | 0.15±0.05/0.03±0.04 | 0.56±0.03/0.07±0.03 |
| | Original | - | 0.13±0.05/0.01±0.02 | 0.15±0.05/0.03±0.04 | 0.56±0.03/0.07±0.03 |
| | *CluSim* | | | | |
| | Metalabels | 1.00±0.00 | 0.87±0.06 | 0.84±0.05 | 0.20±0.05 |
| | Original | - | 0.87±0.06 | 0.84±0.05 | 0.20±0.05 |
| | *Adjusted Rand Index* | | | | |
| | Metalabels | 1.00±0.00 | 0.90±0.05 | 0.88±0.04 | 0.24±0.06 |
| | Original | - | 0.90±0.05 | 0.88±0.04 | 0.24±0.06 |

**Table C. '*Low*' connectivity synthetic network measures of module similarity.** Values for 10 randomly generated network realizations; random subgraphs based on 100 instances. See also Fig B.

| | | Original | Metric | Threshold | Random |
|---|---|---|---|---|---|
| | $m$ | | | | |
| | Metalabels | 4 | - | - | - |
| | Louvain | 4.3±0.5 | 6.1±1.6 | 6.5±1.8 | 16±2 |
| | Infomap | 9.1±2 | 19±3.6 | 16±2.7 | 34±3 |
| Louvain | $y_{AB}$ | | | | |
| | Metalabels | 0.95±0.05 | 0.81±0.11 | 0.78±0.10 | 0.48±0.03 |
| | Original | - | 0.84±0.10 | 0.80±0.10 | 0.47±0.03 |
| | $J_{A\to B}/J_{B\to A}$ | | | | |
| | Metalabels | 0.96±0.02/0.91±0.10 | 0.87±0.08/0.67±0.19 | 0.86±0.09/0.63±0.17 | 0.39±0.05/0.22±0.03 |
| | Original | - | 0.90±0.08/0.72±0.17 | 0.88±0.11/0.66±0.15 | 0.38±0.06/0.22±0.03 |
| | $h_{A\to B}/h_{B\to A}$ | | | | |
| | Metalabels | 0.06±0.04/0.06±0.04 | 0.17±0.08/0.07±0.03 | 0.18±0.08/0.06±0.02 | 0.54±0.04/0.20±0.05 |
| | Original | - | 0.12±0.07/0.03±0.03 | 0.12±0.07/0.03±0.03 | 0.53±0.04/0.21±0.05 |
| | *CluSim* | | | | |
| | Metalabels | 0.95±0.03 | 0.82±0.10 | 0.81±0.10 | 0.29±0.04 |
| | Original | - | 0.86±0.10 | 0.84±0.10 | 0.28±0.04 |
| | *Adjusted Rand Index* | | | | |
| | Metalabels | 0.95±0.03 | 0.85±0.08 | 0.84±0.08 | 0.31±0.05 |
| | Original | - | 0.89±0.07 | 0.88±0.08 | 0.31±0.06 |
| Infomap | $y_{AB}$ | | | | |
| | Metalabels | 0.66±0.07 | 0.46±0.04 | 0.49±0.04 | 0.34±0.01 |
| | Original | - | 0.60±0.04 | 0.64±0.08 | 0.38±0.03 |
| | $J_{A\to B}/J_{B\to A}$ | | | | |
| | Metalabels | 0.96±0.02/0.91±0.10 | 0.87±0.08/0.67±0.19 | 0.86±0.09/0.63±0.17 | 0.22±0.03/0.11±0.01 |
| | Original | - | 0.90±0.08/0.72±0.17 | 0.88±0.11/0.66±0.15 | 0.28±0.05/0.15±0.02 |
| | $h_{A\to B}/h_{B\to A}$ | | | | |
| | Metalabels | 0.06±0.04/0.06±0.04 | 0.17±0.08/0.07±0.03 | 0.18±0.08/0.06±0.02 | 0.62±0.02/0.12±0.03 |
| | Original | - | 0.12±0.07/0.03±0.03 | 0.12±0.07/0.03±0.03 | 0.43±0.05/0.15±0.03 |
| | *CluSim* | | | | |
| | Metalabels | 0.95±0.03 | 0.82±0.10 | 0.81±0.10 | 0.13±0.02 |
| | Original | - | 0.86±0.10 | 0.84±0.10 | 0.16±0.02 |
| | *Adjusted Rand Index* | | | | |
| | Metalabels | 0.95±0.03 | 0.85±0.08 | 0.84±0.08 | 0.15±0.03 |
| | Original | - | 0.89±0.07 | 0.88±0.08 | 0.15±0.03 |

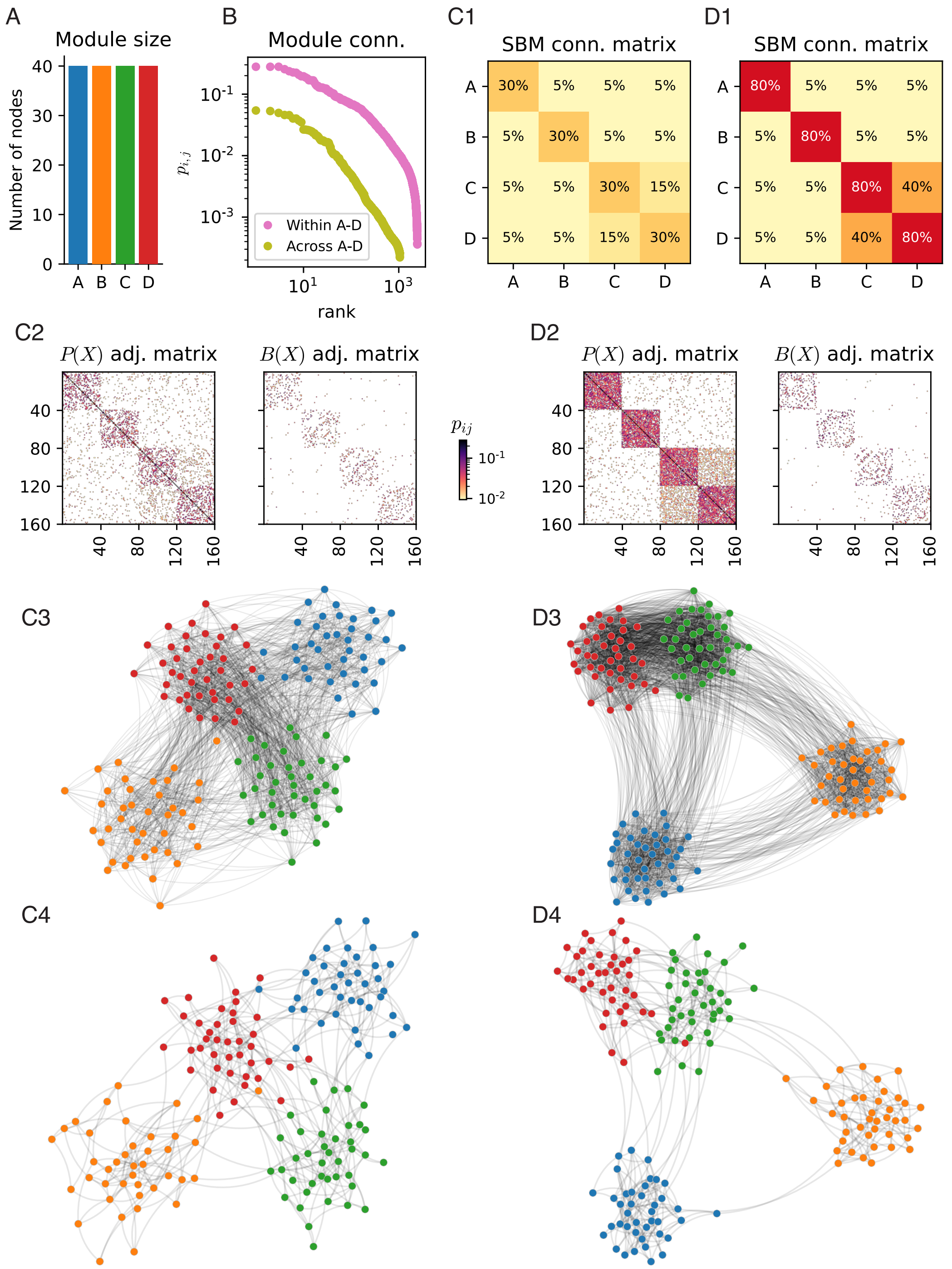


**Fig B. Synthetic networks with connectivity generated by Stochastic Block Models (SBM) and edge weights sampled from the SocioPatterns French Primary School (Fr-PS) contact network.** **A**. Synthetic module sizes. **B**. Proximity values ($p_{ij}$), sampled with replacement from the Fr-PS contact network, used to connect edges within or across modules. **C1 & D1** Two SBM probability matrices are used to generate networks with '*low*' and '*high*' connectivity, respectively. Note the hierarchical structure of modules C and D in both realizations. Two realizations shown, one with '*low*' (C2-C4) and another with '*high*' (D2-D4) module connectivity. **C2-4**. '*Low*' connectivity synthetic network realization. **D2-4**. '*High*' connectivity synthetic network realization. **C2 & D2**. The adjacency matrix of original graph and its metric backbone. **C3 & D3**. Force layout visualization of the original graph, as computed by the NetworkX python package [52]. Node positioning obtained after 100 iterations. **C4 & D4**. Force layout visualization of the metric backbone. Node positioning seeded from the original graph (C3 & C4) and obtained after 100 iterations.

### C Datasets

This section details the social contact datasets upon which we computed our backbones and measures, as well as details of backbone analysis not shown in main manuscript. Sections are arranged by scientific group and project. We thank the authors for making their datasets freely available to the scientific community. These include the works comprised in the SocioPatterns collaboration [37], and references [33] and [34].

#### C.1 SocioPatterns French Primary School (Fr-PS)

This data set is comprised of 125,773 records of contact between 242 individuals (232 students and 10 teachers) in a primary school in Lyon, France. The age of students (elementary cycle) ranges between 6 and 12 years old. Data were collected on October 1$^{st}$ and 2$^{nd}$ in 2009. The school was composed by 5 grades (1$^{st}$ through 5$^{th}$), each of them comprising two classes, for a total of 10 classes. Each class has an assigned room and an assigned teacher. The smallest class has 22 children and the largest 26. The school day runs from 8:30am to 4:30pm, with a lunch break from 12pm to 2pm, and two breaks of 20-25 minutes around 10:30am and 3:30pm. Lunches are served in a common canteen, and a shared playground is located outside the main building. As the playground and the canteen do not have enough capacity to host all the students at the same time, only two or three classes have breaks at the same time, and lunches are taken in two consecutive turns [45]. Additional metadata contains individual roles: teacher, or in the case of students, the class to which they belong. All teachers and 95% of the student population participated in the experiment.

Due to the nature of the social environment in which this data was collected, it is expected that students and their teacher to represent a module. The metalabels of the data, which assign students to their classes should closely represent the natural modular structure of the social environment, which includes the teacher of each class. This is visually depicted in Fig C.

Similarly to other datasets from the SocioPatterns collaboration [37], face-to-face contacts were recorded using active RFID devices, embedded in unobtrusive wearable badges. The badges exchange multi-channel bi-directional radio communication. Their use of low-power signals, which is shielded by the human body, enables the reliable measure of face-to-face proximity, within a distance of approximately 1-1.5 meters, and with temporal resolution of 20 seconds. Details on how this technology is used to monitor social interactions and to identify contact patterns are available in [2, 32, 43].

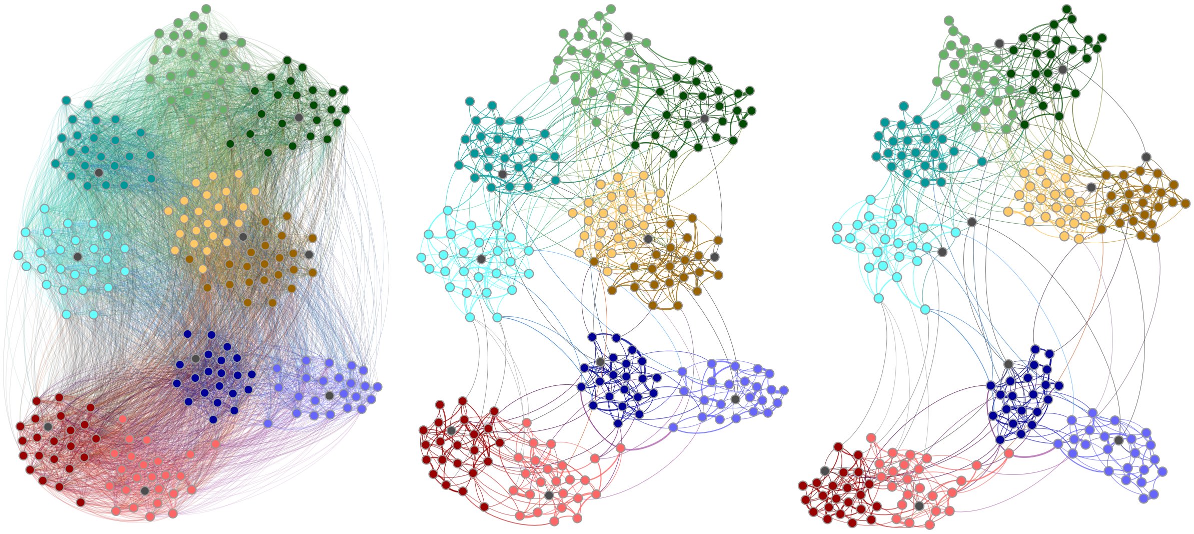

**Fig C. Contacts in the French Primary School (Fr-PS) *social* network.** Distance network, $D(X)$, shown left; backbone, $B(X)$, shown middle and right. Node layout algorithm computed for the original distance network (left and middle) and then recomputed for the backbone (right). Colors represent student grade and class: 1$^{st}$ grades in cyan, 2$^{nd}$ in green, 3$^{rd}$ in orange, 4$^{th}$ in blue, and 5$^{th}$ in red; lighter and darker shades of the same color separate classes within grade; teachers are shown in gray.

**Table D. French Primary School (Fr-PS) metric backbone statistics.**

| | *social* | *individual time* | *experiment time* |
|---|---|---|---|
| $D(X)$ | | | |
| Nodes | 242 | 242 | 242 |
| Edges | 8,317 | 8,317 | 8,317 |
| $B(X)$ | | | |
| Metric edges ($s_{ij} = 1$) | 790 (9.5%) | 764 (9.19%) | 762 (9.16%) |
| Semi-metric edges ($s_{ij} > 1$) | 7,527 (90.5%) | 7,553 (90.81%) | 7,555 (90.84%) |

**Table E. French Primary School (Fr-PS) measures of module similarity.** Values for random subgraphs based on 100 instances.

| | | Original | Metric | Threshold | Random |
|---|---|---|---|---|---|
| | $m$ | | | | |
| | Metalabels | 11 (10*) | - | - | - |
| | Louvain | 8 | 8 | 9 | 21±2.5 |
| | Infomap | 8 | 21 | 19 | 50±3.6 |
| Louvain | $y_{AB}$ | | | | |
| Louvain | Metalabels | 0.89 | 0.86 | 0.85 | 0.53±0.03 |
| Louvain | Original | - | 0.83 | 0.83 | 0.51±0.03 |
| Louvain | $J_{A\to B}/J_{B\to A}$ | | | | |
| Louvain | Metalabels | 0.79/0.87 | 0.75/0.82 | 0.77/0.80 | 0.41±0.05/0.27±0.03 |
| Louvain | Original | - | 0.75/0.76 | 0.78/0.74 | 0.41±0.05/0.25±0.03 |
| Louvain | $h_{A\to B}/h_{B\to A}$ | | | | |
| Louvain | Metalabels | 0.01/0.08 | 0.03/0.11 | 0.05/0.10 | 0.42±0.03/0.25±0.04 |
| Louvain | Original | - | 0.08/0.08 | 0.09/0.06 | 0.45±0.04/0.22±0.04 |
| Louvain | *CluSim* | | | | |
| Louvain | Metalabels | 0.79 | 0.74 | 0.77 | 0.35±0.04 |
| Louvain | Original | - | 0.72 | 0.76 | 0.32±0.04 |
| Louvain | *Adjusted Rand Index* | | | | |
| Louvain | Metalabels | 0.80 | 0.76 | 0.78 | 0.34±0.05 |
| Louvain | Original | - | 0.77 | 0.79 | 0.34±0.06 |
| Infomap | $y_{AB}$ | | | | |
| Infomap | Metalabels | 0.88 | 0.67 | 0.65 | 0.41±0.01 |
| Infomap | Original | - | 0.60 | 0.64 | 0.37±0.01 |
| Infomap | $J_{A\to B}/J_{B\to A}$ | | | | |
| Infomap | Metalabels | 0.78/0.86 | 0.78/0.45 | 0.71/0.44 | 0.27±0.03/0.15±0.01 |
| Infomap | Original | - | 0.78/0.36 | 0.81/0.41 | 0.25±0.03/0.12±0.01 |
| Infomap | $h_{A\to B}/h_{B\to A}$ | | | | |
| Infomap | Metalabels | 0.01/0.09 | 0.16/0.03 | 0.16/0.05 | 0.49±0.02/0.16±0.03 |
| Infomap | Original | - | 0.18/0.02 | 0.16/0.03 | 0.52±0.02/0.14±0.03 |
| Infomap | *CluSim* | | | | |
| Infomap | Metalabels | 0.77 | 0.73 | 0.67 | 0.18±0.02 |
| Infomap | Original | - | 0.63 | 0.72 | 0.15±0.02 |
| Infomap | *Adjusted Rand Index* | | | | |
| Infomap | Metalabels | 0.78 | 0.76 | 0.73 | 0.20±0.03 |
| Infomap | Original | - | 0.61 | 0.74 | 0.16±0.02 |

* when teachers are assigned to their class module.

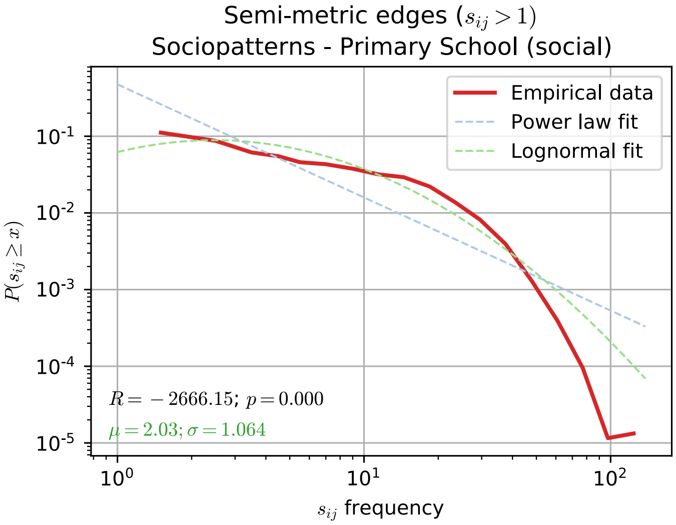


**Fig D. Distribution of semi-metric distortion values in the French Primary School (Fr-PS) network.** Log-binned distribution of semi-metric distortion values ($s_{ij} > 1$) for the $\sigma = 90.5\%$ of semi-metric edges in the French Primary School (Fr-PS) network. Both a log-normal ($\langle s_{ij} \rangle = 2.03$; SD=1.064) and a power law fit are shown; a comparison between the two favours the former as a better representation of the data. Data fitted using the 'powerlaw' python package [3, 4].

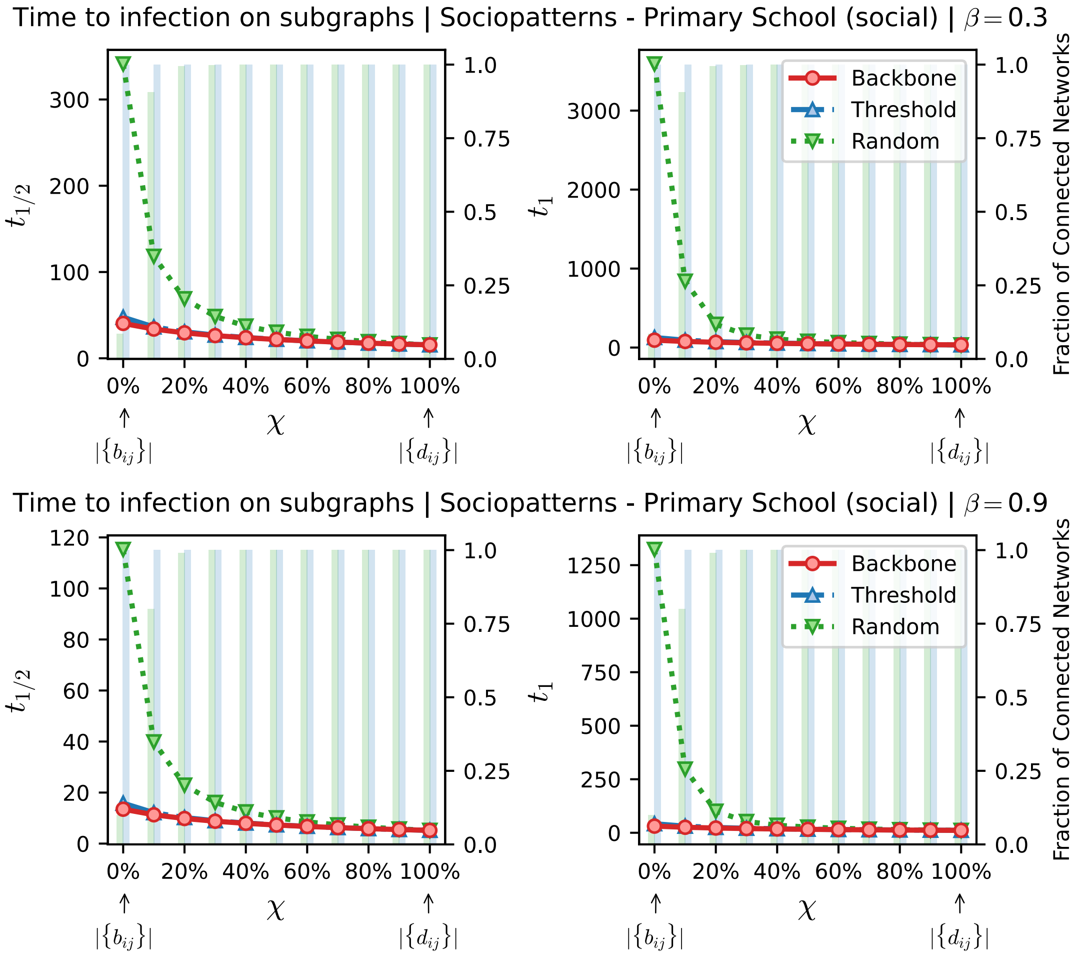


**Fig E. Time to infection using the metric backbone, threshold, or random subgraphs of the French Primary school (Fr-PS) network.** The horizontal axis denotes $\chi$, a parameter to sweep the proportion of edges of the original network that are included in the subgraphs analyzed. When $\chi = 0\%$ (leftmost value on axis) we have the metric backbone subgraph, or threshold and random subgraphs with the same number of edges as the metric backbone (i.e. $|\{b_{ij}\}| = \tau(D).|\{d_{ij}\}|$, per eq. 6). As $\chi$ increases, edges from the original network that are not on the backbone or same-size threshold and random subgraphs, are progressively added until the original network itself is reached at $\chi = 100\%$. (Left panels) denotes the time for half of the population to be infected, $t_{1/2}$. (Right panels) denotes the time for all nodes in the network to be infected, $t_1$. The green and blue bars, quantified against the right vertical axis in each panel, denote the fraction of networks in threshold and random baseline ensembles that are connected for a given $\chi$. Non-connected networks are discarded to compute the spreading times. For the simulations shown, the spreading parameter was set as $\beta = 0.3/p_{max}$ (top panels) or $\beta = 0.9/p_{max}$ (bottom panels) where $p_{max}$ is the largest proximity weight of the original network.

#### C.2 SocioPatterns French High School (Fr-HS)

This data set was gathered over a period of 4 days in 2013. It contains 188,508 contact records between 327 high school students of specific classes called “classes préparatoires”. The data collection took place in Lycée Thiers, Marseilles, France. These classes are specific to the French educational system, taken by the students for two years after the end of the usual high school studies. Since this student body is largely separated from other high school students (i.e., different building and lunch), they constitute an almost closed population with few contacts with the outside world, at least during workdays. These classes are intended to prepare the students to undergo competitive exams yielding admission to various higher education colleges [46]. Student are categorized in classes with three types of specializations: Three “MP” classes with focus on mathematics and physics; two “PC” classes on physics and chemistry; one “PSI” class on engineering; and three “BIO” classes with a focus on biology. The total students participation in the study was 86.3%.

Similar to the Primary School (Section C.1), we also expect the social environment in which the data was collected to be highly modular with respect to the classes and specialization in which the students are enrolled. The metadata, that categorize students in their specialization, and further in each assigned class, should naturally represent a hierarchical order of social affinity. This can be visually seen in Fig F.

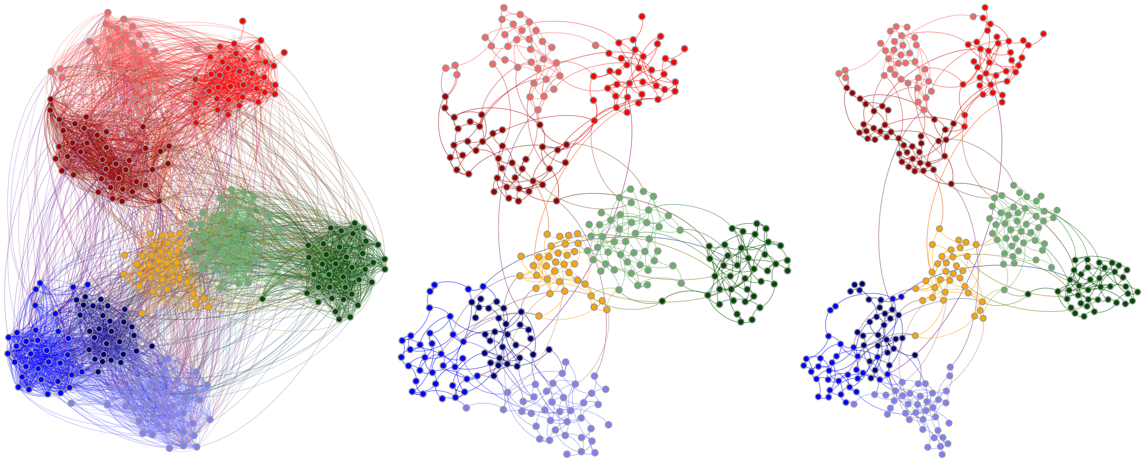

**Fig F. Contacts in the French High School (Fr-HS) *social* network.** Distance network, $D(X)$, shown left; backbone, $B(X)$, shown middle and right. Node layout algorithm computed for the original distance network (left and middle) and then recomputed for the backbone (right). Colors represent the student specialization: “MP” in blue, “PC” in green, “PSI” in orange, and “BIO” in red; lighter or darker colors separate classes within specialization.

**Table F. French High School (Fr-HS) measures of module similarity.** Values for random subgraphs based on 100 instances.

| | | Original | Metric | Threshold | Random |
|---|---|---|---|---|---|
| | $m$ | | | | |
| | Metalabels | 9 | - | - | - |
| | Louvain | 10 | 17 | 27 | 48±3.8 |
| | Infomap | 36 | 66 | 66 | 88±3.9 |
| Louvain | $y_{AB}$ | | | | |
| | Metalabels | 0.88 | 0.68 | 0.55 | 0.38±0.02 |
| | Original | - | 0.74 | 0.61 | 0.37±0.02 |
| | $J_{A\to B}/J_{B\to A}$ | | | | |
| | Metalabels | 0.89/0.81 | 0.70/0.46 | 0.68/0.30 | 0.36±0.05/0.14±0.01 |
| | Original | - | 0.77/0.55 | 0.76/0.37 | 0.34±0.05/0.14±0.01 |
| | $h_{A\to B}/h_{B\to A}$ | | | | |
| | Metalabels | 0.09/0.08 | 0.24/0.11 | 0.25/0.07 | 0.44±0.03/0.12±0.02 |
| | Original | - | 0.17/0.05 | 0.18/0.01 | 0.41±0.03/0.12±0.02 |
| | *CluSim* | | | | |
| | Metalabels | 0.87 | 0.62 | 0.57 | 0.27±0.04 |
| | Original | - | 0.67 | 0.63 | 0.26±0.04 |
| | *Adjusted Rand Index* | | | | |
| | Metalabels | 0.89 | 0.71 | 0.68 | 0.30±0.04 |
| | Original | - | 0.76 | 0.75 | 0.29±0.04 |
| Infomap | $y_{AB}$ | | | | |
| | Metalabels | 0.48 | 0.37 | 0.37 | 0.31±0.01 |
| | Original | - | 0.64 | 0.68 | 0.41±0.01 |
| | $J_{A\to B}/J_{B\to A}$ | | | | |
| | Metalabels | 0.62/0.23 | 0.27/0.13 | 0.34/0.13 | 0.20±0.02/0.09±0.01 |
| | Original | - | 0.69/0.46 | 0.78/0.49 | 0.28±0.02/0.18±0.01 |
| | $h_{A\to B}/h_{B\to A}$ | | | | |
| | Metalabels | 0.29/0.03 | 0.46/0.03 | 0.44/0.03 | 0.53±0.01/0.09±0.01 |
| | Original | - | 0.11/0.04 | 0.09/0.03 | 0.28±0.01/0.14±0.01 |
| | *CluSim* | | | | |
| | Metalabels | 0.49 | 0.17 | 0.21 | 0.11±0.01 |
| | Original | - | 0.41 | 0.49 | 0.17±0.01 |
| | *Adjusted Rand Index* | | | | |
| | Metalabels | 0.58 | 0.23 | 0.28 | 0.14±0.01 |
| | Original | - | 0.35 | 0.43 | 0.13±0.01 |

**Table G. French High School (Fr-HS) metric backbone statistics.**

| | *social* | *individual time* | *experiment time* |
|---|---|---|---|
| $D(X)$ | | | |
| Nodes | 327 | 327 | 327 |
| Edges | 5,818 | 5,818 | 5,818 |
| $B(X)$ | | | |
| Metric edges ($s_{ij} = 1$) | 603 (10.36%) | 590 (10.14%) | 600 (10.31%) |
| Semi-metric edges ($s_{ij} > 1$) | 5,215 (90.10%) | 5,228 (89.86%) | 5,218 (89.69%) |

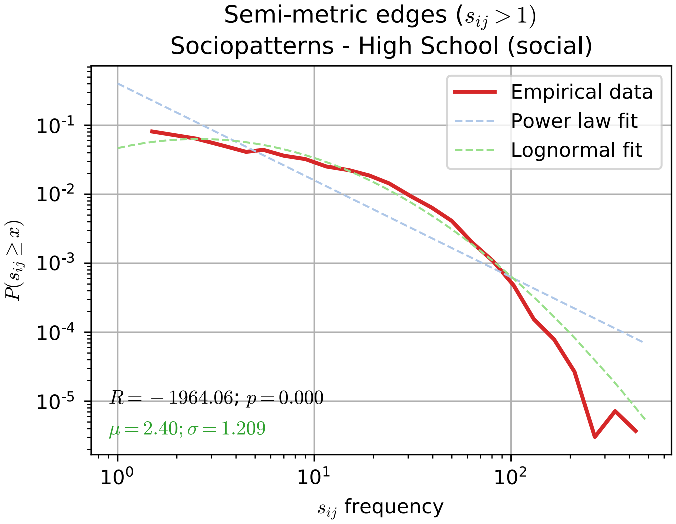


**Fig G. Distribution of semi-metric distortion values in the French High School (Fr-HS) network.** Log-binned distribution of semi-metric distortion values ($s_{ij} > 1$) for the $\sigma = 90.10\%$ of semi-metric edges in the French High School (Fr-HS) network. Both a log-normal ($\langle s_{ij} \rangle = 2.40$; SD=1.209) and a power law fit are shown; a comparison between the two favours the former as a better representation of the data. Data fitted using the 'powerlaw' python package [3, 4].

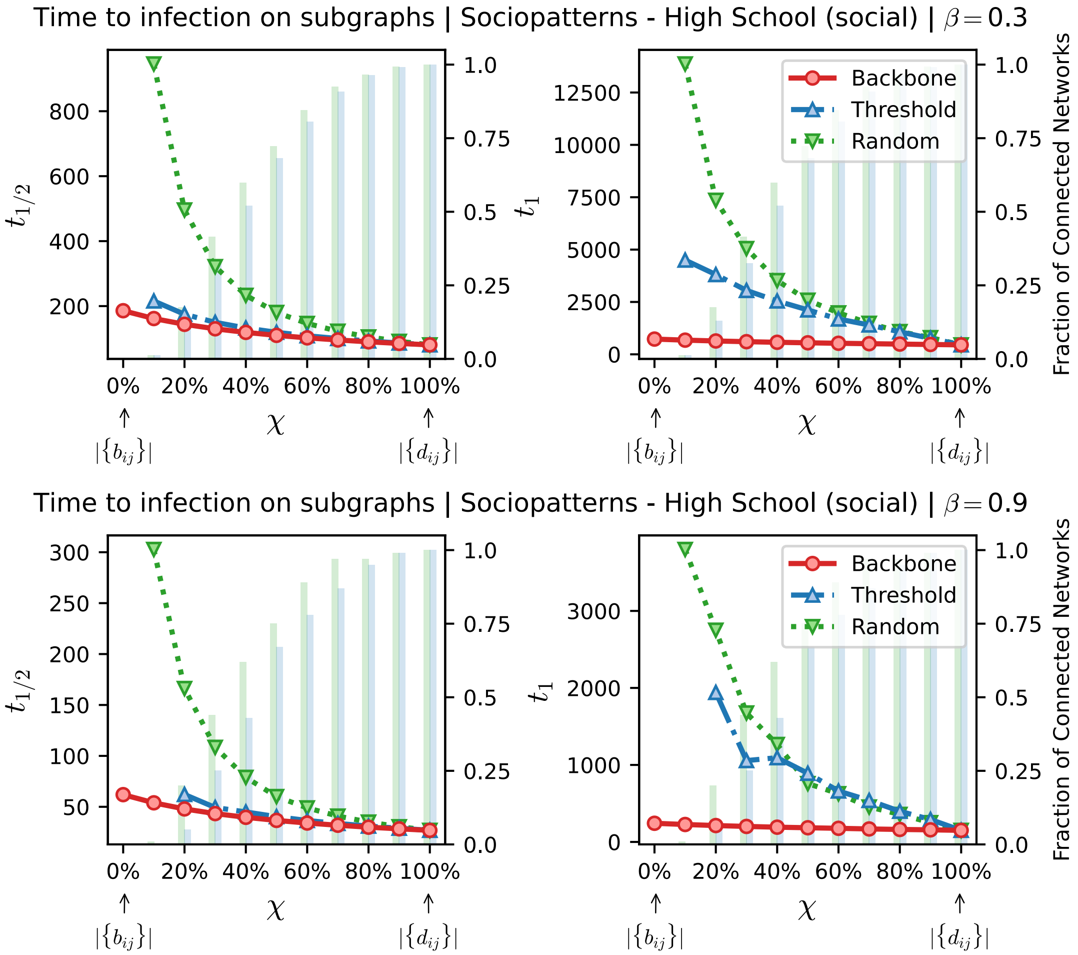


**Fig H. Time to infection using the metric backbone, threshold, or random subgraphs of the French High School (Fr-HS) network.** The horizontal axis denotes $\chi$, a parameter to sweep the proportion of edges of the original network that are included in the subgraphs analyzed. When $\chi = 0\%$ (leftmost value on axis) we have the metric backbone subgraph, or threshold and random subgraphs with the same number of edges as the metric backbone (i.e. $|\{b_{ij}\}| = \tau(D).|\{d_{ij}\}|$, per eq. 6). As $\chi$ increases, edges from the original network that are not on the backbone or same-size threshold and random subgraphs, are progressively added until the original network itself is reached at $\chi = 100\%$. (Left panels) denotes the time for half of the population to be infected, $t_{1/2}$. (Right panels) denotes the time for all nodes in the network to be infected, $t_1$. The green and blue bars, quantified against the right vertical axis in each panel, denote the fraction of networks in threshold and random baseline ensembles that are connected for a given $\chi$. Non-connected networks are discarded to compute the spreading times. For the simulations shown, the spreading parameter was set as $\beta = 0.3/p_{max}$ (top panels) or $\beta = 0.9/p_{max}$ (bottom panels) where $p_{max}$ is the largest proximity weight of the original network.

#### C.3 SocioPatterns ACM Hypertext 2009 Scientific Conference (It-SC)

This data set was gathered over a period of 3 days in 2009. It contains 20,818 contact records between 113 individuals attending the ACM Hypertext 2009 Conference in Torino, Italy [43]. The proportion of participants that volunteered was 75%. No additional metadata was recorded and, due to the nature of the social environment in which the data were collected, we do not expect individuals to form any highly modular structure (see Fig I), especially since scientists attend conferences to promote their research to the widest possible audience. Thus, there is a social incentive to meet and network with the widest possible number of individuals. Having said that, it could still be possible to disentangle (likely weak) modules from the data, if additional knowledge were known about the participants. For instance, their scientific seniority, advisor-advisee relationships, who attended particular workshops or parallel sessions during the conference, etc. However, none of this additional social information is available in the metadata.

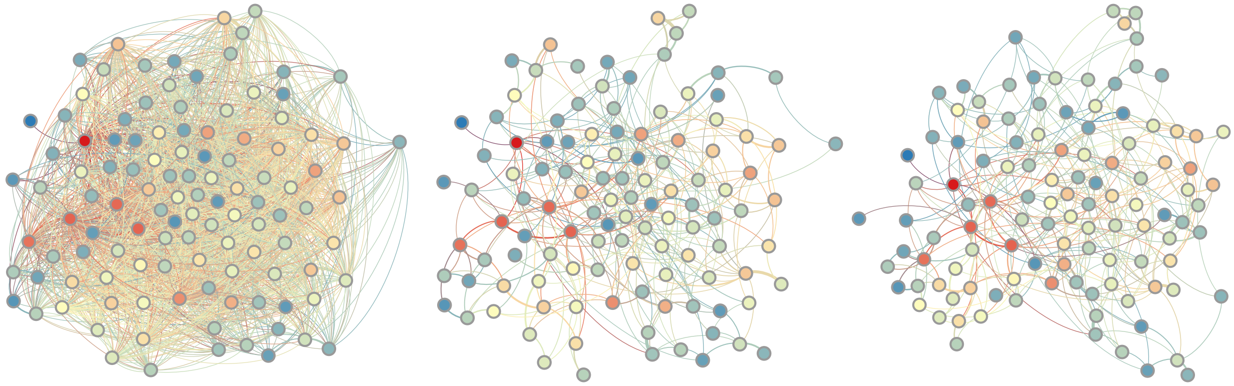

**Fig I. Contacts in the Italian Scientific Conference (It-SC) *social* network.** Distance network, $D(X)$, shown left; backbone, $B(X)$, shown middle and right. Node layout algorithm computed for the original distance network (left and middle) and then recomputed for the backbone (right). Colors represent node degree.

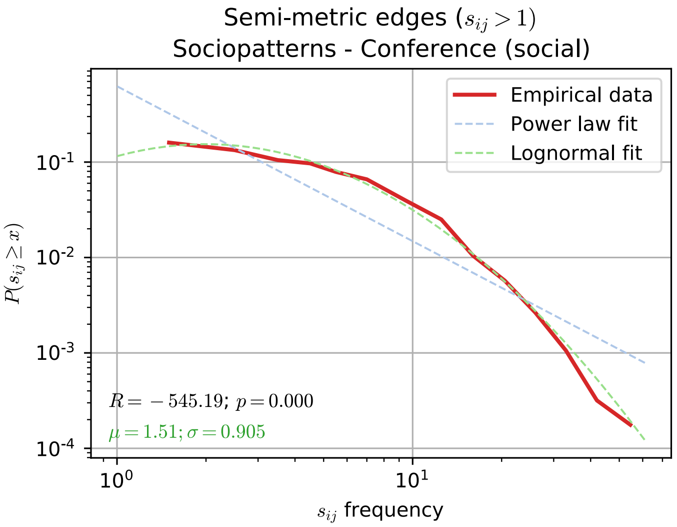


**Fig J. Distribution of semi-metric distortion values in the Italian Scientific Conference (It-SC) network.** Log-binned distribution of semi-metric distortion values ($s_{ij} > 1$) for the $\sigma = 85.97\%$ of semi-metric edges in the Italian Scientific Conference (It-SC) network. Both a log-normal ($\langle s_{ij} \rangle = 1.51$; SD=0.905) and a power law fit are shown; a comparison between the two favours the former as a better representation of the data. Data fitted using the 'powerlaw' python package [3, 4].

**Table H. Italian Scientific Conference (It-SC) measures of module similarity.** Values for random subgraphs based on 100 instances.

| | | Original | Metric | Threshold | Random |
|---|---|---|---|---|---|
| | $m$ | | | | |
| | Louvain | 10 | 11 | 14 | 16±2.1 |
| | Infomap | 8 | 21 | 24 | 28±2.8 |
| Louvain | $y_{AB}$ | | | | |
| | Original | - | 0.81 | 0.72 | 0.45±0.03 |
| | $J_{A\to B}/J_{B\to A}$ | | | | |
| | Original | - | 0.74/0.70 | 0.74/0.56 | 0.22±0.03/0.18±0.02 |
| | $h_{A\to B}/h_{B\to A}$ | | | | |
| | Original | - | 0.16/0.14 | 0.14/0.08 | 0.55±0.03/0.39±0.06 |
| | *CluSim* | | | | |
| | Original | - | 0.65 | 0.68 | 0.18±0.02 |
| | *Adjusted Rand Index* | | | | |
| | Original | - | 0.61 | 0.70 | 0.05±0.02 |
| Infomap | $y_{AB}$ | | | | |
| | Original | - | 0.56 | 0.50 | 0.36±0.02 |
| | $J_{A\to B}/J_{B\to A}$ | | | | |
| | Original | - | 0.79/0.34 | 0.72/0.28 | 0.29±0.05/0.14±0.02 |
| | $h_{A\to B}/h_{B\to A}$ | | | | |
| | Original | - | 0.11/0.04 | 0.14/0.04 | 0.42±0.03/0.18±0.03 |
| | *CluSim* | | | | |
| | Original | - | 0.32 | 0.30 | 0.13±0.01 |
| | *Adjusted Rand Index* | | | | |
| | Original | - | 0.07 | 0.07 | 0.01±0.01 |

**Table I. Italian Scientific Conference (It-SC) metric backbone statistics.**

| | *social* | *individual time* | *experiment time* |
|---|---|---|---|
| $D(X)$ | | | |
| Nodes | 113 | 113 | 113 |
| Edges | 2,196 | 2,196 | 2,196 |
| $B(X)$ | | | |
| Metric edges ($s_{ij} = 1$) | 308 (14.03%) | 305 (13.89%) | 253 (11.52%) |
| Semi-metric edges ($s_{ij} > 1$) | 1,888 (85.97%) | 1,891 (86.11%) | 1,943 (88.48%) |

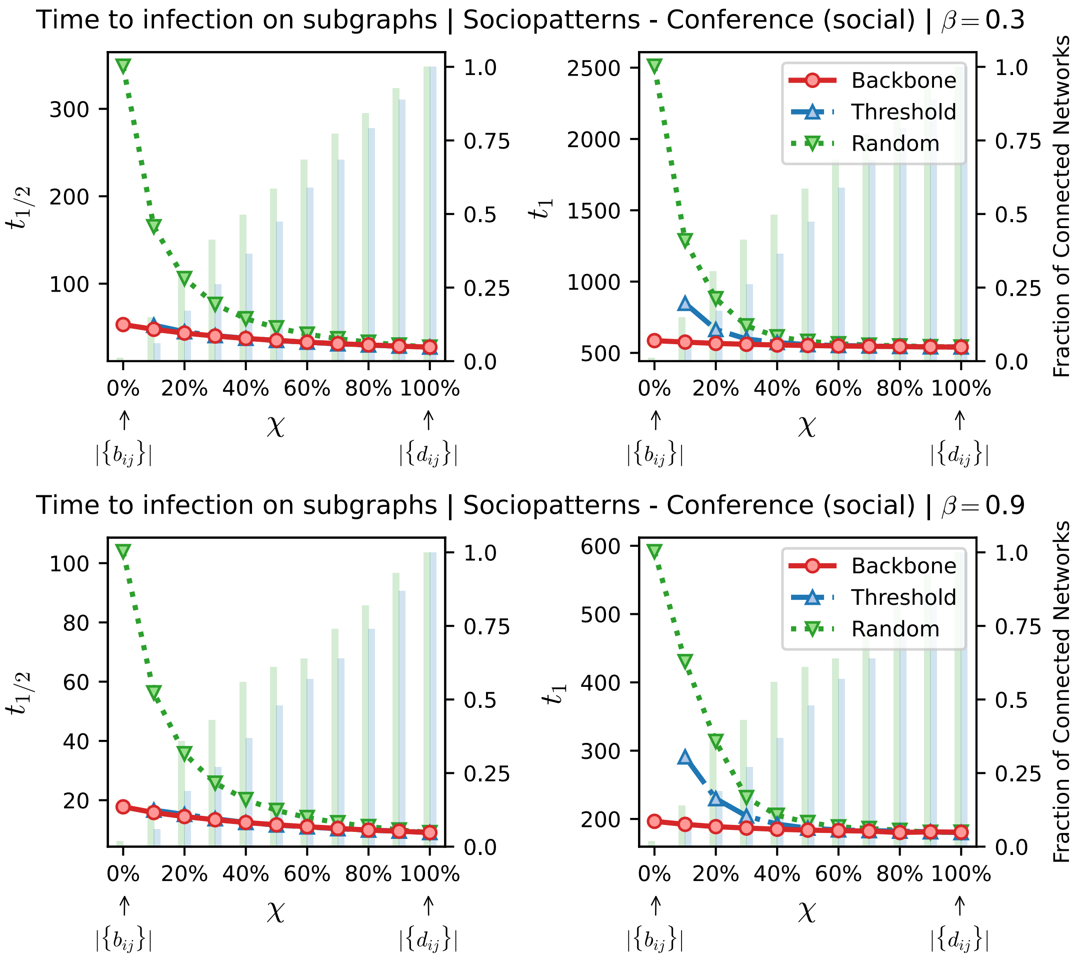


**Fig K. Time to infection using the metric backbone, threshold, or random subgraphs of the Italian Scientific Conference (It-SC) network.** The horizontal axis denotes $\chi$, a parameter to sweep the proportion of edges of the original network that are included in the subgraphs analyzed. When $\chi = 0\%$ (leftmost value on axis) we have the metric backbone subgraph, or threshold and random subgraphs with the same number of edges as the metric backbone (i.e. $|\{b_{ij}\}| = \tau(D).|\{d_{ij}\}|$, per eq. 6). As $\chi$ increases, edges from the original network that are not on the backbone or same-size threshold and random subgraphs, are progressively added until the original network itself is reached at $\chi = 100\%$. (Left panels) denotes the time for half of the population to be infected, $t_{1/2}$. (Right panels) denotes the time for all nodes in the network to be infected, $t_1$. The green and blue bars, quantified against the right vertical axis in each panel, denote the fraction of networks in threshold and random baseline ensembles that are connected for a given $\chi$. Non-connected networks are discarded to compute the spreading times. For the simulations shown, the spreading parameter was set as $\beta = 0.3/p_{max}$ (top panels) or $\beta = 0.9/p_{max}$ (bottom panels) where $p_{max}$ is the largest proximity weight of the original network.

#### C.4 SocioPatterns Geriatric Ward of French Hospital (Fr-Ho)

This data set was gathered over a period of 4 days, from December 6$^{\text{th}}$ to the 10$^{\text{th}}$, 2010. The data set contains 32,424 contact records between 46 hospital employees and 29 patients (75 total individuals). It was collected in a short stay geriatric unit (19 beds) of a university hospital of almost 1,000 beds, located in Lyon, France [42]. During the collection period, 50 professional staff worked in the unit and 31 patients were present (92% participation rate).

Each day 2 teams of 2 nurses and 3 nurses' aides worked in the ward: one of the teams was present from 7am to 1:30pm and the other from 1:30pm to 8pm. An additional nurse and an additional nurse' aid were moreover present from 9am to 5pm. Two nurses were present during the nights from 8pm to 7am. In addition, a physiotherapist and a nutritionist were present each day at various points in time, with no fixed schedule, and a social counselor and a physical therapist visited on demand (in our analysis they are considered as nurses). Two physicians and 2 interns were present from 8:00 am to 17:00 pm each day. Visits were allowed from 12am to 8pm but visitors were not included in the study. Metadata denote the roles of individuals, such as patient, nurse, medical doctor, or administrative staff.

Due to the nature of the environment in which the data were collected we expect the roles of individuals not to be particularly meaningful in regards to any underlying modular social structure. Perhaps as expected, patients are scattered through the network (see red nodes in Fig L), and surrounded by nurses (orange nodes). Most doctors (blue nodes) are strongly connected in the proximity network as well as in the backbone, denoting they spent much of experiment in close proximity to each other, irrespective of whether they were interns or not. Based also on the network visual characterization, it is also likely the administrative staff (green nodes) had widely different roles among themselves. One green node is highly central while the majority is at the periphery of the network.

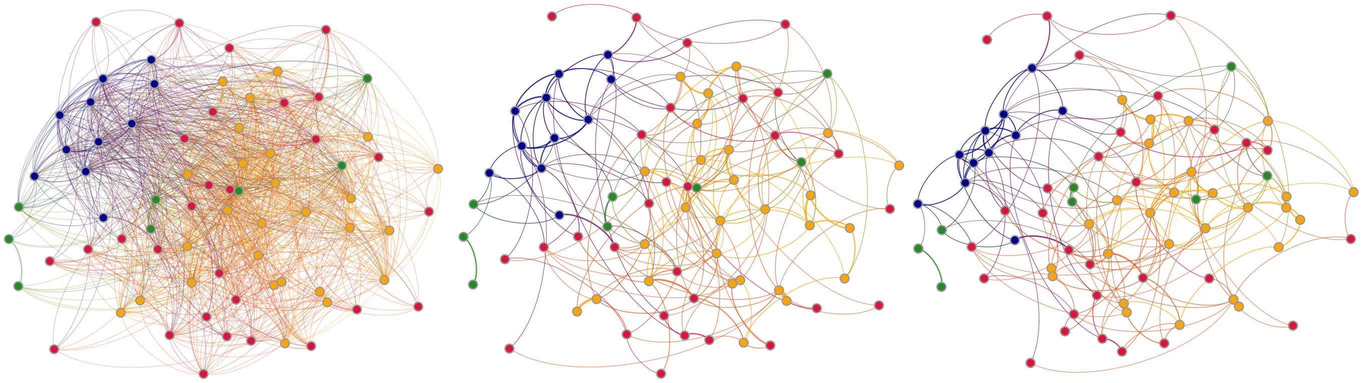

**Fig L. Contacts in the French Hospital (Fr-Ho) *social* network.** Distance network, $D(X)$, shown left; backbone, $B(X)$, shown middle and right. Node layout algorithm computed for the original distance network (left and middle) and then recomputed for the backbone (right). Colors represent the role of individuals: patients in red, nurses in orange, medical doctors in blue, and staff in green; lighter or darker colors separate classes within specialization.

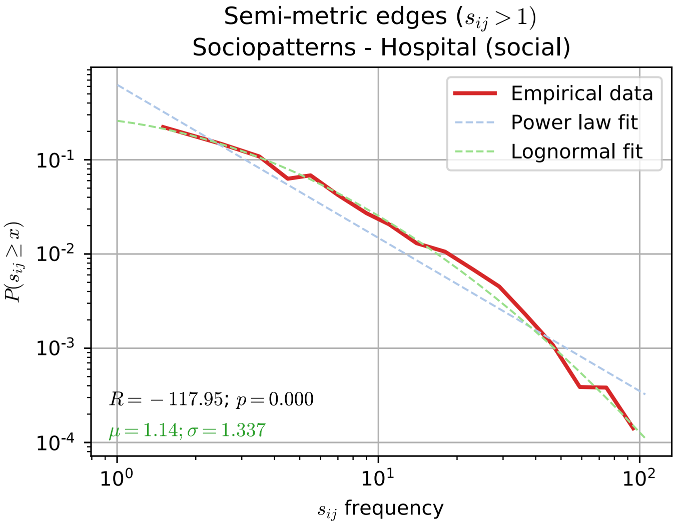


**Fig M. Distribution of semi-metric distortion values in the French Hospital (Fr-Ho) network.** Log-binned distribution of semi-metric distortion values ($s_{ij} > 1$) for the $\sigma = 80.95\%$ of semi-metric edges in the French Hospital (Fr-Ho) network. Both a log-normal ($\langle s_{ij} \rangle = 1.14$; SD=1.337) and a power law fit are shown; a comparison between the two favours the former as a better representation of the data. Data fitted using the 'powerlaw' python package [3, 4].

**Table J. French Hospital (Fr-Ho) measures of module similarity.** Values for random subgraphs based on 100 instances.

| | | Original | Metric | Threshold | Random |
|---|---|---|---|---|---|
| | $m$ | | | | |
| | Metalabels | 4 | - | - | - |
| | Louvain | 9 | 10 | 15 | 11±1.7 |
| | Infomap | 10 | 13 | 16 | 16±2.3 |
| Louvain | $y_{AB}$ | | | | |
| | Metalabels | 0.48 | 0.46 | 0.39 | 0.44±0.03 |
| | Original | - | 0.88 | 0.73 | 0.51±0.04 |
| | $J_{A\to B}/J_{B\to A}$ | | | | |
| | Metalabels | 0.33/0.21 | 0.30/0.19 | 0.30/0.14 | 0.24±0.03/0.16±0.02 |
| | Original | - | 0.85/0.81 | 0.81/0.55 | 0.29±0.06/0.27±0.05 |
| | $h_{A\to B}/h_{B\to A}$ | | | | |
| | Metalabels | 0.61/0.47 | 0.61/0.42 | 0.55/0.28 | 0.68±0.05/0.49±0.09 |
| | Original | - | 0.11/0.04 | 0.16/0.04 | 0.47±0.05/0.37±0.07 |
| | *CluSim* | | | | |
| | Metalabels | 0.25 | 0.22 | 0.21 | 0.19±0.03 |
| | Original | - | 0.74 | 0.67 | 0.25±0.03 |
| | *Adjusted Rand Index* | | | | |
| | Metalabels | 0.09 | 0.09 | 0.08 | 0.04±0.03 |
| | Original | - | 0.72 | 0.68 | 0.11±0.04 |
| Infomap | $y_{AB}$ | | | | |
| | Metalabels | 0.45 | 0.44 | 0.39 | 0.40±0.02 |
| | Original | - | 0.83 | 0.65 | 0.47±0.04 |
| | $J_{A\to B}/J_{B\to A}$ | | | | |
| | Metalabels | 0.38/0.21 | 0.30/0.17 | 0.30/0.14 | 0.21±0.04/0.12±0.01 |
| | Original | - | 0.87/0.72 | 0.70/0.49 | 0.31±0.07/0.25±0.04 |
| | $h_{A\to B}/h_{B\to A}$ | | | | |
| | Metalabels | 0.49/0.37 | 0.58/0.39 | 0.54/0.29 | 0.69±0.04/0.48±0.06 |
| | Original | - | 0.08/0.05 | 0.14/0.08 | 0.38±0.04/0.27±0.05 |
| | *CluSim* | | | | |
| | Metalabels | 0.33 | 0.21 | 0.21 | 0.15±0.03 |
| | Original | - | 0.62 | 0.51 | 0.22±0.03 |
| | *Adjusted Rand Index* | | | | |
| | Metalabels | 0.16 | 0.09 | 0.09 | 0.02±0.02 |
| | Original | - | 0.46 | 0.37 | 0.10±0.04 |

**Table K. French Hospital (Fr-Ho) metric backbone statistics.**

| | *social* | *individual time* | *experiment time* |
|---|---|---|---|
| $D(X)$ | | | |
| Nodes | 75 | 75 | 75 |
| Edges | 1,139 | 1,139 | 1,139 |
| $B(X)$ | | | |
| Metric edges ($s_{ij} = 1$) | 217 (19.05%) | 204 (17.91%) | 161 (14.14%) |
| Semi-metric edges ($s_{ij} > 1$) | 922 (80.95%) | 935 (82.09%) | 978 (85.86%) |

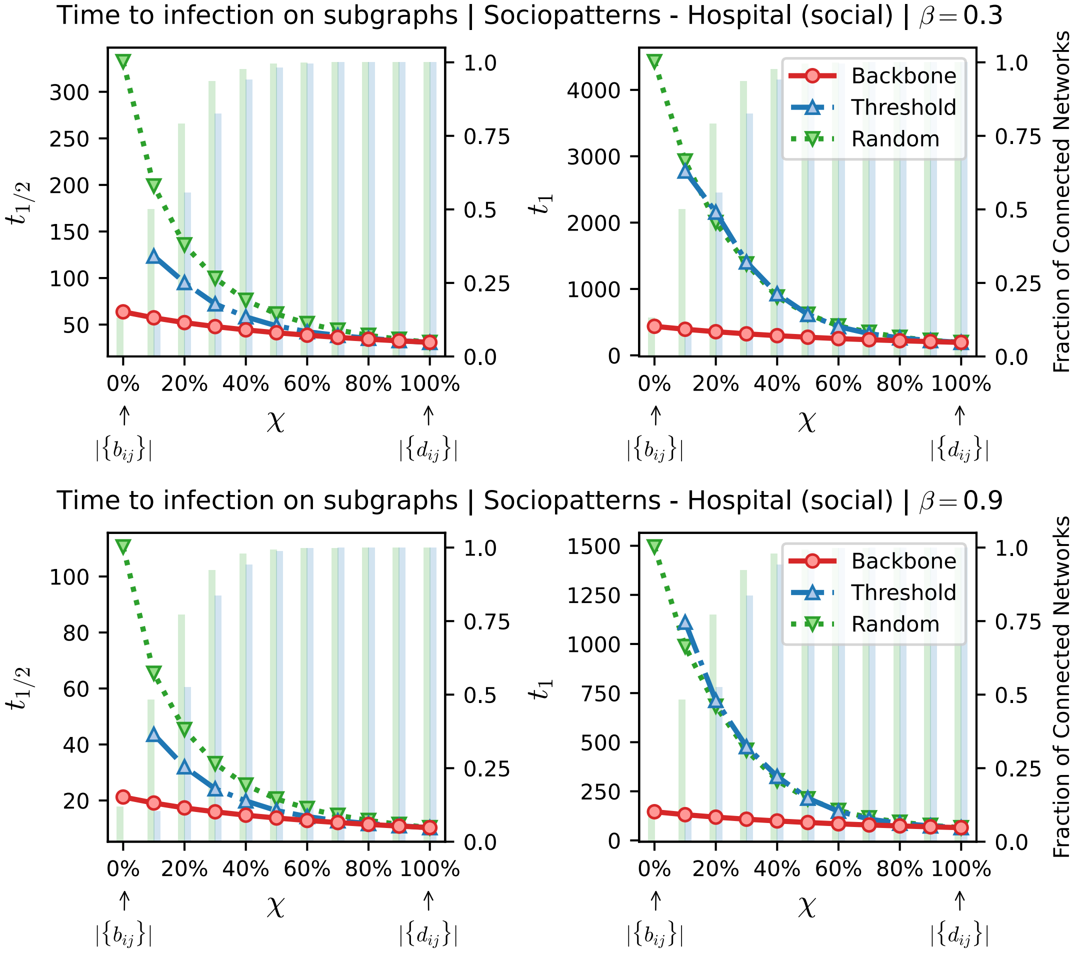


**Fig N. Time to infection using the metric backbone, threshold, or random subgraphs of the French Hospital (Fr-Ho) network.** The horizontal axis denotes $\chi$, a parameter to sweep the proportion of edges of the original network that are included in the subgraphs analyzed. When $\chi = 0\%$ (leftmost value on axis) we have the metric backbone subgraph, or threshold and random subgraphs with the same number of edges as the metric backbone (i.e. $|\{b_{ij}\}| = \tau(D).|\{d_{ij}\}|$, per eq. 6). As $\chi$ increases, edges from the original network that are not on the backbone or same-size threshold and random subgraphs, are progressively added until the original network itself is reached at $\chi = 100\%$. (Left panels) denotes the time for half of the population to be infected, $t_{1/2}$. (Right panels) denotes the time for all nodes in the network to be infected, $t_1$. The green and blue bars, quantified against the right vertical axis in each panel, denote the fraction of networks in threshold and random baseline ensembles that are connected for a given $\chi$. Non-connected networks are discarded to compute the spreading times. For the simulations shown, the spreading parameter was set as $\beta = 0.3/p_{max}$ (top panels) or $\beta = 0.9/p_{max}$ (bottom panels) where $p_{max}$ is the largest proximity weight of the original network.

#### C.5 SocioPatterns Contacts in a Workplace (Fr-Wo)

This data set is comprised of 78,249 contact records between 232 individuals working in one of the two office buildings of the French Institute for Public Health Surveillance (*Institut de veille sanitaire*; InVS), located near Paris, France. Data collection lasted 12 days in 2015 [44]. Additional metadata contained the departments in which individuals worked. The 12 different departments were of different sizes ranging from 60 to 2 individuals. For instance, the Scientific Direction (DISQ), the Department of Chronic Diseases and Traumatisms (DMCT), the Department of Health and Environment (DSE), Human Resources (SRH), Logistics (SFLE), or other unspecified departments, such as DMI, DST, DCAR, SSI, SCOM, SDOC or DG. DMI is the largest department with 60 individuals. In second, with little more then half, is DSE with 34 individuals, followed by DMCT with 33, DST with 24, DISQ with 18, SFLE with 17, and DCAR with 15. Departments with less than ten

**Table L. French Workplace (Fr-Wo) metric backbone statistics.**

| | *social* | *individual time* | *experiment time* |
|---|---|---|---|
| $D(X)$ | | | |
| Nodes (isolates) | 232 (15) | 232 (15) | 232 (15) |
| Edges | 4,274 | 4,274 | 4,274 |
| $B(X)$ | | | |
| Metric edges ($s_{ij} = 1$) | 745 (17.43%) | 712 (16.66%) | 708 (16.57%) |
| Semi-metric edges ($s_{ij} > 1$) | 3,529 (82.57%) | 3,562 (83.34%) | 3,566 (83.43%) |

individuals were SSI and SRH with 9, SCOM with 7, SDOC with 4, and lastly DG with 2. Some of the nodes in the network were isolates (i.e. had no connection to another node in the network), therefore in our analysis and plots we used only the network largest connected component.

The social environment of a workplace is naturally conducive of a hierarchical and modular organization. Employees working on the same projects or affiliated to the same department are expected to form socially coherent and modular structures. Conversely, employees holding management or overseeing positions are expected to navigate one or more these modules, thus being more central in the network (see Fig O). Therefore, we expect the metalabels in this data set, that describe each of the departments, to be a coherent characterization of the underlying social structure of this workplace.

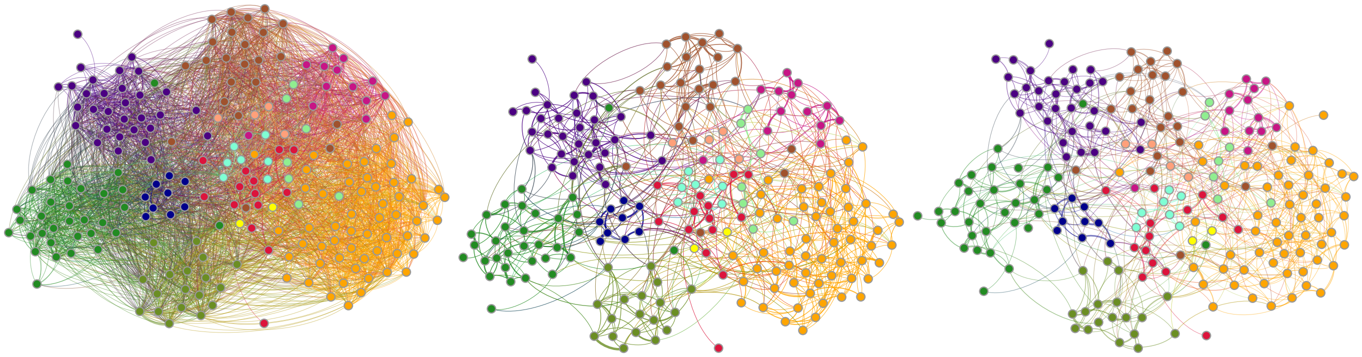

**Fig O. Contacts in the French Workplace (Fr-Wo) *social* network.** Distance network, $D(X)$, shown left; backbone, $B(X)$, shown middle and right. Node layout algorithm computed for the original distance network (left and middle) and then recomputed for the backbone (right). Colors represent different departments. DMI (orange), DSE (dark green), DMCT (purple), DST (brown), DISQ (olive), SFLE (crimson red), DCAR (violet), SSI (lime green), SRH (dark blue), SCOM (aquamarine), SDOC (light salmon), DG (yellow). Only the largest connected component is shown.

**Table M. French Workplace (Fr-Wo) measures of module similarity.** Values for random subgraphs based on 100 instances. Isolates were removed for computations.

| | | Original | Metric | Threshold | Random |
|---|---|---|---|---|---|
| | $m$ | | | | |
| | Metalabels | 12 | - | - | - |
| | Louvain | 10 | 11 | 17 | 21±2.1 |
| | Infomap | 16 | 27 | 30 | 46±2.8 |
| Louvain | $y_{AB}$ | | | | |
| | Metalabels | 0.87 | 0.78 | 0.60 | 0.45±0.02 |
| | Original | - | 0.85 | 0.64 | 0.46±0.02 |
| | $J_{A\to B}/J_{B\to A}$ | | | | |
| | Metalabels | 0.79/0.91 | 0.68/0.75 | 0.59/0.45 | 0.32±0.05/0.23±0.03 |
| | Original | - | 0.82/0.78 | 0.68/0.44 | 0.35±0.05/0.22±0.03 |
| | $h_{A\to B}/h_{B\to A}$ | | | | |
| | Metalabels | 0.08/0.07 | 0.15/0.14 | 0.17/0.10 | 0.41±0.04/0.27±0.03 |
| | Original | - | 0.12/0.10 | 0.17/0.09 | 0.45±0.04/0.27±0.04 |
| | *CluSim* | | | | |
| | Metalabels | 0.87 | 0.67 | 0.56 | 0.28±0.04 |
| | Original | - | 0.71 | 0.57 | 0.28±0.04 |
| | *Adjusted Rand Index* | | | | |
| | Metalabels | 0.89 | 0.63 | 0.55 | 0.26±0.05 |
| | Original | - | 0.68 | 0.54 | 0.25±0.05 |
| Infomap | $y_{AB}$ | | | | |
| | Metalabels | 0.73 | 0.60 | 0.57 | 0.41±0.02 |
| | Original | - | 0.70 | 0.67 | 0.44±0.02 |
| | $J_{A\to B}/J_{B\to A}$ | | | | |
| | Metalabels | 0.73/0.61 | 0.68/0.38 | 0.70/0.34 | 0.16±0.04/0.16±0.01 |
| | Original | - | 0.81/0.52 | 0.80/0.47 | 0.32±0.04/0.19±0.02 |
| | $h_{A\to B}/h_{B\to A}$ | | | | |
| | Metalabels | 0.12/0.05 | 0.21/0.10 | 0.20/0.06 | 0.43±0.02/0.18±0.02 |
| | Original | - | 0.12/0.04 | 0.11/0.04 | 0.38±0.02/0.19±0.02 |
| | *CluSim* | | | | |
| | Metalabels | 0.65 | 0.47 | 0.51 | 0.18±0.02 |
| | Original | - | 0.57 | 0.56 | 0.19±0.02 |
| | *Adjusted Rand Index* | | | | |
| | Metalabels | 0.69 | 0.42 | 0.48 | 0.15±0.03 |
| | Original | - | 0.41 | 0.45 | 0.12±0.02 |

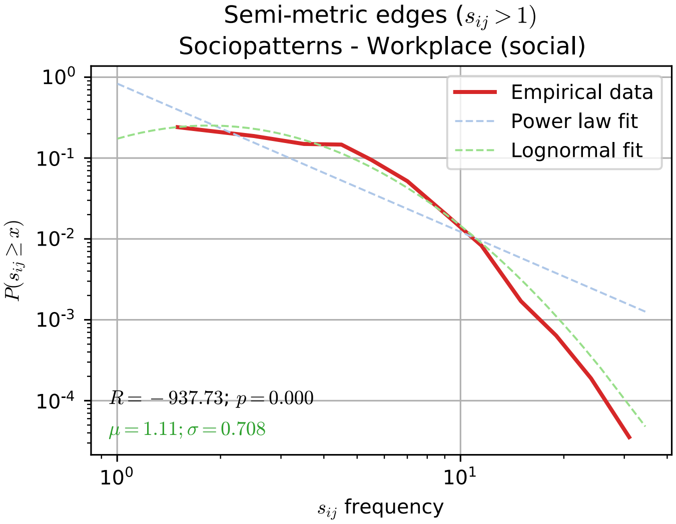


**Fig P. Distribution of semi-metric distortion values in the French Workplace (Fr-Wo) network.** Log-binned distribution of semi-metric distortion values ($s_{ij} > 1$) for the $\sigma = 82.57.5\%$ of semi-metric edges in the French Workplace (Fr-Wo) network. Both a log-normal ($\langle s_{ij} \rangle = 1.11$; SD=0.708) and a power law fit are shown; a comparison between the two favours the former as a better representation of the data. Data fitted using the 'powerlaw' python package [3, 4].

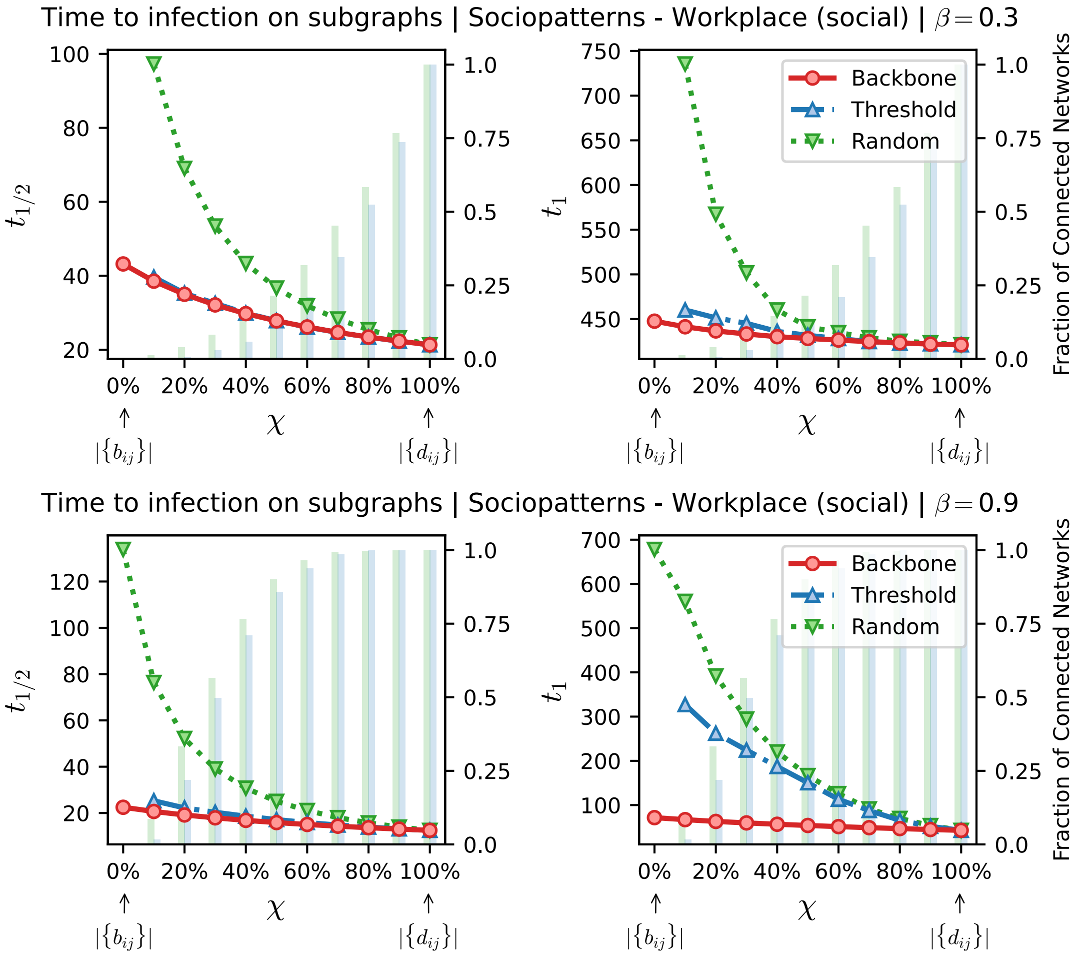


**Fig Q. Time to infection using the metric backbone, threshold, or random subgraphs of the French Workplace (Fr-Wo) network.** The horizontal axis denotes $\chi$, a parameter to sweep the proportion of edges of the original network that are included in the subgraphs analyzed. When $\chi = 0\%$ (leftmost value on axis) we have the metric backbone subgraph, or threshold and random subgraphs with the same number of edges as the metric backbone (i.e. $|\{b_{ij}\}| = \tau(D).|\{d_{ij}\}|$, per eq. 6). As $\chi$ increases, edges from the original network that are not on the backbone or same-size threshold and random subgraphs, are progressively added until the original network itself is reached at $\chi = 100\%$. (Left panels) denotes the time for half of the population to be infected, $t_{1/2}$. (Right panels) denotes the time for all nodes in the network to be infected, $t_1$. The green and blue bars, quantified against the right vertical axis in each panel, denote the fraction of networks in threshold and random baseline ensembles that are connected for a given $\chi$. Non-connected networks are discarded to compute the spreading times. For the simulations shown, the spreading parameter was set as $\beta = 0.3/p_{max}$ (top panels) or $\beta = 0.9/p_{max}$ (bottom panels) where $p_{max}$ is the largest proximity weight of the original network.

#### C.6 SocioPatterns Stay Away Exhibit at Dublin Science Gallery (Ir-Ex)

This data set consists of social contacts collected from visitors to the *Infectious: Stay Away* exhibit at the Science Gallery in Dublin, Ireland. The exhibit ran for about three months, attracting more than 14,000 visitors with more than 230,000 face-to-face contacts recorded [43]. Upon entering the exhibit, visitors received an RFID tag, which enabled the successful tracking of almost the totality of contacts. Since this was an art exhibit, visitors spent little time at the experiment in comparison to their overall museum time, or even with the total time the museum was open to public. Visitors also followed a pre-defined path, interacting with the exposition at different points and locations. Because of this dynamic, the observed network closely follows the social relationships of groups of individuals—groups of friends or families—interacting with the exhibit through time, which gives the daily networks larger diameter than in other settings. No additional metadata information was supplied. Because of device availability, node unique identifiers were restarted daily, meaning that we must consider an independent aggregated network for each day. For this reason, measures on these networks presented in Table O were averaged over all the 69 days.

In Figure R we show the aggregated network for July 15$^{\text{th}}$ 2009, the day with the highest number of recorded participants.

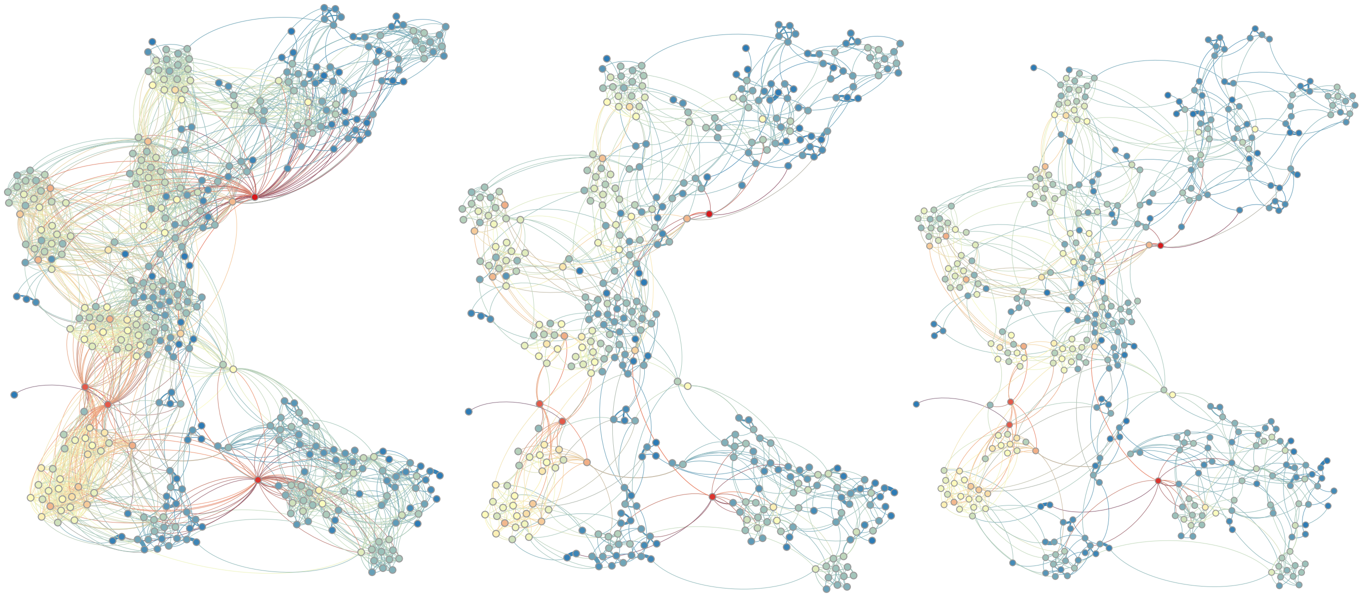

**Fig R. Contacts in the Irish Stay Away Exhibit (Ir-Ex) *social* network for July 15$^{\text{th}}$, 2009.** Distance network, $D(X)$, shown left; backbone, $B(X)$, shown middle and right. Node layout algorithm computed for the original distance network (left and middle) and then recomputed for the backbone (right). Colors denote node degree.

**Table N. Irish Stay Away Exhibit (Ir-Ex) measures of module similarity.** Values for July 15$^{th}$, 2009. Random subgraphs based on 100 instances.

| | | Original | Metric | Threshold | Random |
|---|---|---|---|---|---|
| | $m$ | | | | |
| | Louvain | 30 | 38 | 51 | 54±4 |
| | Infomap | 75 | 79 | 83 | 86±4.6 |
| Louvain | $y_{AB}$ | | | | |
| | Original | - | 0.79 | 0.73 | 0.54±0.03 |
| | $J_{A\to B}/J_{B\to A}$ | | | | |
| | Original | - | 0.79/0.66 | 0.84/0.55 | 0.54±0.04/0.35±0.03 |
| | $h_{A\to B}/h_{B\to A}$ | | | | |
| | Original | - | 0.08/0.04 | 0.09/0.02 | 0.19±0.01/0.08±0.01 |
| | *CluSim* | | | | |
| | Original | - | 0.75 | 0.75 | 0.55±0.03 |
| | *Adjusted Rand Index* | | | | |
| | Original | - | 0.82 | 0.82 | 0.64±0.03 |
| Infomap | $y_{AB}$ | | | | |
| | Original | - | 0.95 | 0.90 | 0.66±0.03 |
| | $J_{A\to B}/J_{B\to A}$ | | | | |
| | Original | - | 0.95/0.92 | 0.91/0.84 | 0.57±0.04/0.53±0.03 |
| | $h_{A\to B}/h_{B\to A}$ | | | | |
| | Original | - | 0.01/0.01 | 0.09/0.02 | 0.11±0.01/0.07±0.01 |
| | *CluSim* | | | | |
| | Original | - | 0.92 | 0.88 | 0.58±0.03 |
| | *Adjusted Rand Index* | | | | |
| | Original | - | 0.92 | 0.90 | 0.66±0.05 |

**Table O. Irish Stay Away Exhibit (Ir-Ex) metric backbone statistics.** Differently from other datasets, numbers shown here represent the average (± std.) value for all the 69 days the exhibit was measured.

| | *social* | *individual time* | *experiment time* |
|---|---|---|---|
| $D(X)$ | | | |
| Nodes | 159±63 | 159±63 | 159±63 |
| Edges | 645±468 | 645±468 | 645±468 |
| $B(X)$ | | | |
| Metric edges ($s_{ij}=1$) | 283±166 (48±9%) | 272±158 (47±9%) | 321±190 (55±12%) |
| Semi-metric edges ($s_{ij}>1$) | 362±306 (3±1%) | 373±314 (3±1%) | 325±289 (2±3%) |

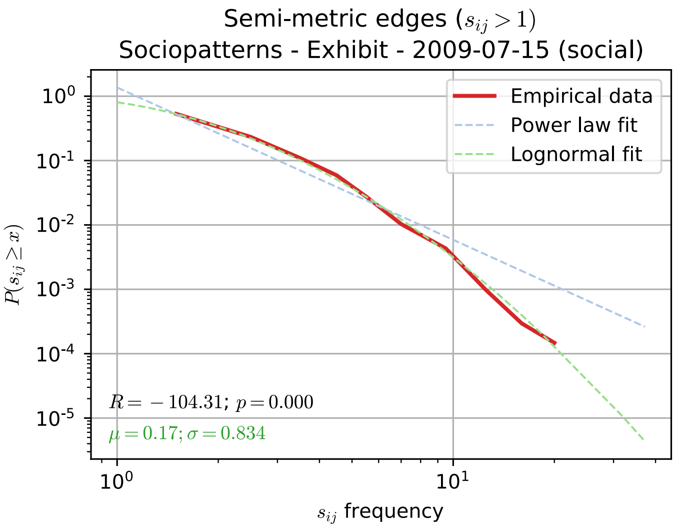


**Fig S. Distribution of semi-metric distortion values in the Irish Stay Away Exhibit (Ir-Ex) network.** Log-binned distribution of semi-metric distortion values ($s_{ij} > 1$) for the $\sigma = 90.10\%$ of semi-metric edges in the Irish Stay Away Exhibit (Ir-Ex) network.Values for July 15$^{\text{th}}$, 2009. Both a log-normal ($\langle s_{ij} \rangle = 0.17$; SD=0.834) and a power law fit are shown; a comparison between the two favours the former as a better representation of the data. Data fitted using the 'powerlaw' python package [3, 4].

#### C.7 Salathé et al. American High School (US-HS)

This data set was gathered on January 14$^{\text{th}}$ 2010, a typical high school day in the United States (unspecified city/state). Data were collected from students, teachers and staff via wireless "sensor network motes" (TelosB; Crossbow Technologies Inc.), with data covering 94% of the entire school population [33]. Metadata for this network differentiates students, teachers, and school staff (see Fig T). Unfortunately, the metadata do not contain student classes and therefore we cannot describe student classes as network modules as it would be expected from the underlying natural social environment. Since the metalabels in this network do not represent the coherent social structure, we have omitted in Table P the module comparison with metalabels. In addition, and differently from the SocioPatterns datasets, the contact data made available by the authors was not grouped by time window but rather by the sum of contact pair occurrences. To our analysis, this means we could only compute the "social" normalization as described in Section A.

**Table P. American High School (US-HS) measures of module similarity.** Values for random subgraphs based on 100 instances.

| | | Original | Metric | Threshold | Random |
|---|---|---|---|---|---|
| | $m$ | | | | |
| | Metalabels | 4 | - | - | - |
| | Louvain | 6 | 7 | 9 | 29±2.5 |
| | Infomap | 11 | 16 | 17 | 120±4.6 |
| Louvain | $y_{AB}$ | | | | |
| | Original | - | 0.87 | 0.75 | 0.38±0.01 |
| | $J_{A \to B}/J_{B \to A}$ | | | | |
| | Original | - | 0.88/0.77 | 0.88/0.77 | 0.23±0.04/0.11±0.01 |
| | $h_{A \to B}/h_{B \to A}$ | | | | |
| | Original | - | 0.16/0.20 | 0.19/0.12 | 0.69±0.03/0.43±0.05 |
| | *CluSim* | | | | |
| | Original | - | 0.81 | 0.81 | 0.13±0.02 |
| | *Adjusted Rand Index* | | | | |
| | Original | - | 0.81 | 0.79 | 0.01±0.01 |
| Infomap | $y_{AB}$ | | | | |
| | Original | - | 0.76 | 0.71 | 0.25±0.01 |
| | $J_{A \to B}/J_{B \to A}$ | | | | |
| | Original | - | 0.80/0.61 | 0.79/0.55 | 0.19±0.03/0.06±0.01 |
| | $h_{A \to B}/h_{B \to A}$ | | | | |
| | Original | - | 0.13/0.13 | 0.11/0.10 | 0.51±0.01/0.23±0.01 |
| | *CluSim* | | | | |
| | Original | - | 0.77 | 0.72 | 0.04±0.01 |
| | *Adjusted Rand Index* | | | | |
| | Original | - | 0.82 | 0.75 | 0.03±0.01 |

**Table Q. American High School (US-HS) metric backbone statistics.**

| | *social* |
|---|---|
| $D(X)$ | |
| Nodes | 788 |
| Edges | 118,291 |
| $B(X)$ | |
| Metric edges ($s_{ij} = 1$) | 9,275 (7.84%) |
| Semi-metric edges ($s_{ij} > 1$) | 109,016 (92.16%) |

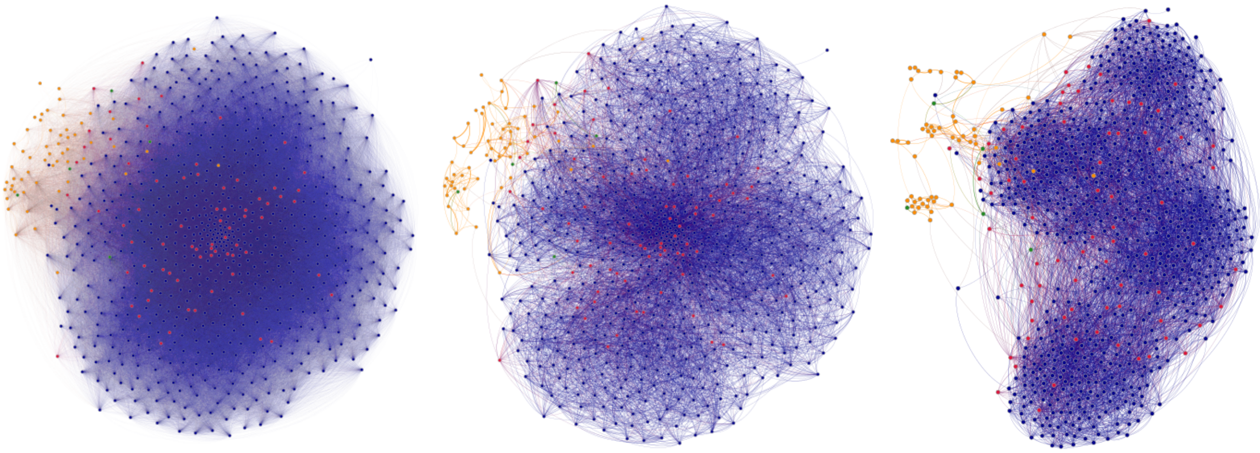

**Fig T. Contacts in the American High-School (US-HS) network.** Distance network, $D(X)$, shown left; backbone, $B(X)$, shown middle and right. Nodes colors denote students in blue, teachers in red, staff in orange. Other (unspecified) individuals are shown in green.

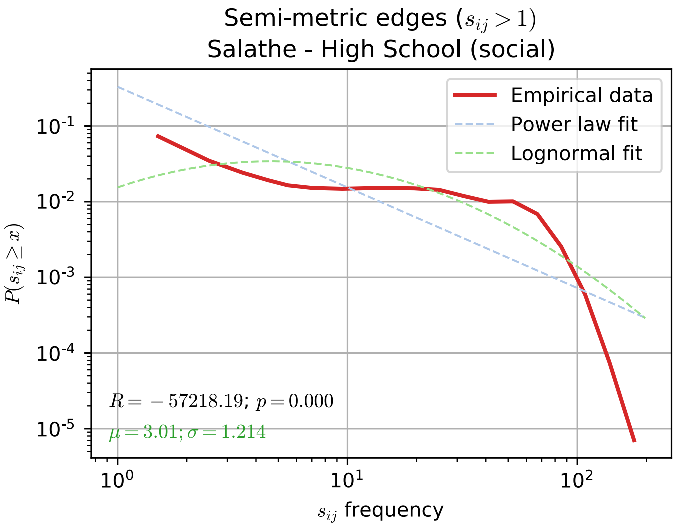


**Fig U. Distribution of semi-metric distortion values in the American High School (US-HS) network.** Log-binned distribution of semi-metric distortion values ($s_{ij} > 1$) for the $\sigma = 92.16\%$ of semi-metric edges in the American High School (US-HS) network. Both a log-normal ($\langle s_{ij} \rangle = 3.01$; SD=1.214) and a power law fit are shown; a comparison between the two favours the former as a better representation of the data. Data fitted using the 'powerlaw' python package [3, 4].

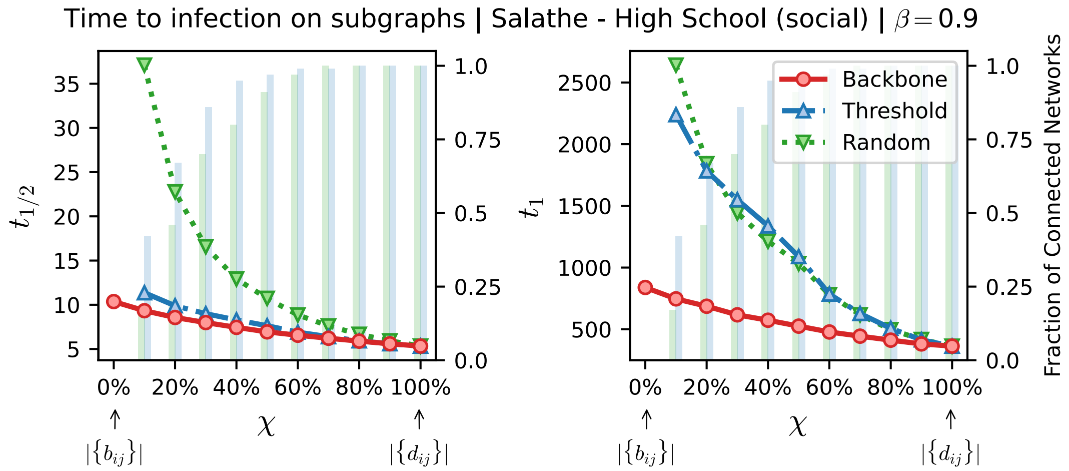


**Fig V. Time to infection using the metric backbone, threshold, or random subgraphs of the American High School (US-HS) network.** The horizontal axis denotes $\chi$, a parameter to sweep the proportion of edges of the original network that are included in the subgraphs analyzed. When $\chi = 0\%$ (leftmost value on axis) we have the metric backbone subgraph, or threshold and random subgraphs with the same number of edges as the metric backbone (i.e. $|\{b_{ij}\}| = \tau(D).|\{d_{ij}\}|$, per eq. 6). As $\chi$ increases, edges from the original network that are not on the backbone or same-size threshold and random subgraphs, are progressively added until the original network itself is reached at $\chi = 100\%$. (Left panels) denotes the time for half of the population to be infected, $t_{1/2}$. (Right panels) denotes the time for all nodes in the network to be infected, $t_1$. The green and blue bars, quantified against the right vertical axis in each panel, denote the fraction of networks in threshold and random baseline ensembles that are connected for a given $\chi$. Non-connected networks are discarded to compute the spreading times. For the simulations shown, the spreading parameter was set as $\beta = 0.3/p_{max}$ (top panels) or $\beta = 0.9/p_{max}$ (bottom panels) where $p_{max}$ is the largest proximity weight of the original network.

#### C.8 Toth et al. American Elementary School (US-SC)

This dataset was gathered in a suburban elementary school in Utah (USA) during two days, January 31$^{th}$ and February 2$^{nd}$, 2013. Metadata consists of gender and grades (K-6); there were 21 different classes across the seven grades. The authors also note that seating arrangements may not have been the same across the two deployment days, as teachers generally changed their classroom seating arrangement at the end of a calendar month. Students from different classes also had the opportunity to mix during lunch, recess, and assembly during the morning of the first deployment day, and a science fair during the afternoon of the second deployment day. School hours were 8:30am to 2:45pm on each day for all students except for kindergartners, for whom school hours were either 8:30am to 11:05am or 12:10 to 2:45pm.

Data availability was similar to the SocioPatterns datasets, but collected using Wireless Ranging Enabled Nodes (WRENs). WRENs were distributed to students in the first-period classrooms and collected before the end of the second day (only data from school hours were made available). Data resolution is approximately 20 seconds (precisely 20,000 binary milliseconds, which equals 19.53125 seconds; there are 210 = 1024 binary milliseconds per second). The processing of the original data can be seen in reference [5]. Three networks were made available in the original work [34], one for each day of collection, and a third containing both days. In our analysis we used the combined network containing both days.

Similar to the other datasets collected in a school environment (see Sections C.1 and C.2, we expect the metalabel classes here to coherently represent the underlying social structure, as can be seen in Fig W. With the students being organized both in grades and classes, we expect the latter to best describe the social structure.

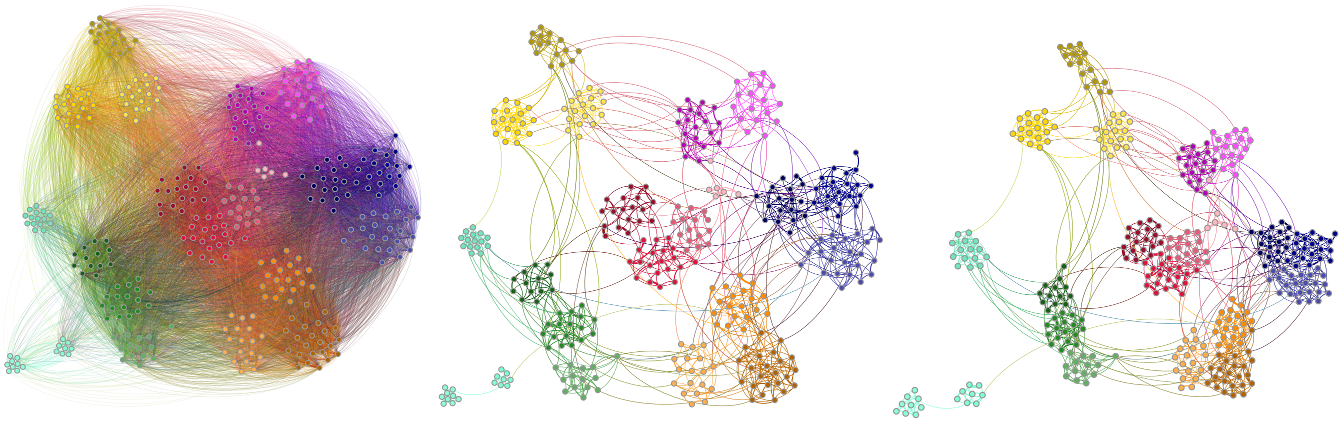

**Fig W. Contacts in the American Elementary School (US-ES).** Distance network, $D(X)$, shown left; backbone, $B(X)$, shown middle and right. Node layout algorithm computed for the original distance network (left and middle) and then recomputed for the backbone (right). Colors denote student grade and class: kindergartners in cyan, 1$^{st}$ grade in yellow, 2$^{nd}$ in green, 3$^{rd}$ in orange, 4$^{th}$ in pink, 5$^{th}$ in blue; and 6$^{th}$ in red; lighter or darker shades of the same color separate classes within grade.

**Table R. American Elementary School (US-ES) measures of module similarity.** Values for random subgraphs based on 100 instances.

| | | Original | Metric | Threshold | Random |
|---|---|---|---|---|---|
| | $m$ | | | | |
| | Metalabels | 21 | - | - | - |
| | Louvain | 15 | 18 | 20 | 38±3.4 |
| | Infomap | 21 | 27 | 25 | 80±4.6 |
| Louvain | $y_{AB}$ | | | | |
| | Metalabels | 0.85 | 0.93 | 0.97 | 0.50±0.02 |
| | Original | - | 0.91 | 0.87 | 0.48±0.02 |
| | $J_{A\to B}/J_{B\to A}$ | | | | |
| | Metalabels | 0.71/0.85 | 0.86/0.94 | 0.95/0.97 | 0.31±0.03/0.23±0.02 |
| | Original | - | 0.91/0.83 | 0.87/0.75 | 0.32±0.03/0.21±0.02 |
| | $h_{A\to B}/h_{B\to A}$ | | | | |
| | Metalabels | 0.00/0.08 | 0.00/0.04 | 0.01/0.01 | 0.41±0.02/0.26±0.03 |
| | Original | - | 0.05/0.00 | 0.07/0.00 | 0.45±0.02/0.24±0.03 |
| | *CluSim* | | | | |
| | Metalabels | 0.71 | 0.86 | 0.94 | 0.26±0.02 |
| | Original | - | 0.83 | 0.74 | 0.24±0.02 |
| | *Adjusted Rand Index* | | | | |
| | Metalabels | 0.70 | 0.86 | 0.93 | 0.24±0.03 |
| | Original | - | 0.84 | 0.75 | 0.24±0.03 |
| Infomap | $y_{AB}$ | | | | |
| | Metalabels | 1.00 | 0.86 | 0.91 | 0.45±0.01 |
| | Original | - | 0.86 | 0.91 | 0.45±0.01 |
| | $J_{A\to B}/J_{B\to A}$ | | | | |
| | Metalabels | 1.00/1.00 | 0.93/0.75 | 0.97/0.84 | 0.27±0.02/0.17±0.01 |
| | Original | - | 0.93/0.75 | 0.97/0.84 | 0.27±0.02/0.17±0.01 |
| | $h_{A\to B}/h_{B\to A}$ | | | | |
| | Metalabels | 0.00/0.00 | 0.05/0.02 | 0.02/0.01 | 0.43±0.01/0.17±0.01 |
| | Original | - | 0.05/0.02 | 0.02/0.01 | 0.43±0.01/0.17±0.01 |
| | *CluSim* | | | | |
| | Metalabels | 1.00 | 0.89 | 0.95 | 0.18±0.02 |
| | Original | - | 0.89 | 0.95 | 0.18±0.02 |
| | *Adjusted Rand Index* | | | | |
| | Metalabels | 1.00 | 0.91 | 0.96 | 0.19±0.02 |
| | Original | - | 0.91 | 0.96 | 0.19±0.02 |

**Table S. American Elementary School (US-ES) metric backbone statistics.**

| | *social* | *individual time* | *experiment time* |
|---|---|---|---|
| $D(X)$ | | | |
| Nodes | 339 | 339 | 339 |
| Edges | 16,546 | 16,546 | 16,546 |
| $B(X)$ | | | |
| Metric edges ($s_{ij} = 1$) | 1,128 (6.82%) | 1,071 (6.47%) | 1,111 (6.71%) |
| Semi-metric edges ($s_{ij} > 1$) | 15,418 (93.18%) | 15,475 (93.53%) | 15,435 (93.29%) |

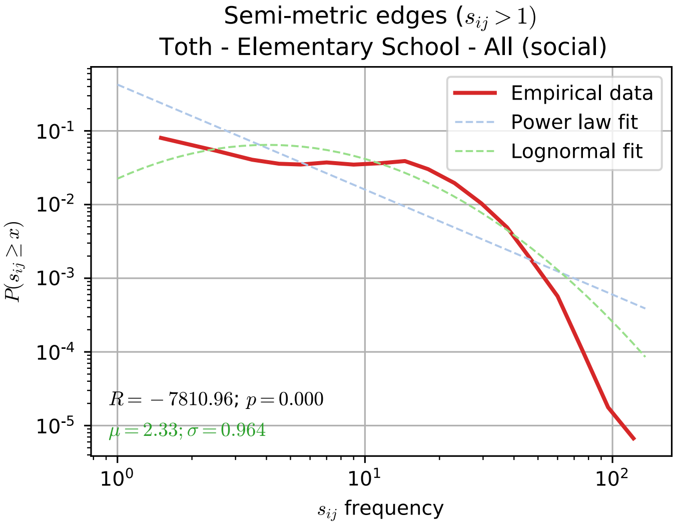


**Fig X. Distribution of semi-metric distortion values in the American Elementary School (US-ES) network.** Log-binned distribution of semi-metric distortion values ($s_{ij} > 1$) for the $\sigma = 93.18\%$ of semi-metric edges in the American Elementary School (US-ES) network. Both a log-normal ($\langle s_{ij} \rangle = 2.33$; SD=0.964) and a power law fit are shown; a comparison between the two favours the former as a better representation of the data. Data fitted using the 'powerlaw' python package [3, 4].

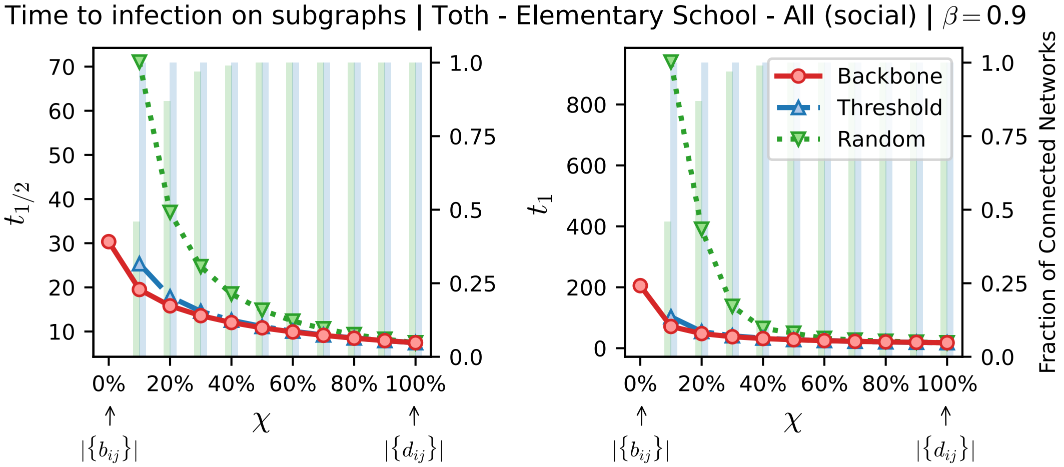


**Fig Y. Time to infection using the metric backbone, threshold, or random subgraphs of the American Elementary School (US-ES) network.** The horizontal axis denotes $\chi$, a parameter to sweep the proportion of edges of the original network that are included in the subgraphs analyzed. When $\chi = 0\%$ (leftmost value on axis) we have the metric backbone subgraph, or threshold and random subgraphs with the same number of edges as the metric backbone (i.e. $|\{b_{ij}\}| = \tau(D).|\{d_{ij}\}|$, per eq. 6). As $\chi$ increases, edges from the original network that are not on the backbone or same-size threshold and random subgraphs, are progressively added until the original network itself is reached at $\chi = 100\%$. (Left panels) denotes the time for half of the population to be infected, $t_{1/2}$. (Right panels) denotes the time for all nodes in the network to be infected, $t_1$. The green and blue bars, quantified against the right vertical axis in each panel, denote the fraction of networks in threshold and random baseline ensembles that are connected for a given $\chi$. Non-connected networks are discarded to compute the spreading times. For the simulations shown, the spreading parameter was set as $\beta = 0.9/p_{max}$ where $p_{max}$ is the largest proximity weight of the original network.

#### C.9 Toth et al. American Middle School (US-MS)

This data set was gathered in a suburban middle school in Utah (USA) during two days, November 28$^{th}$ and 29$^{th}$, 2012. Metadata consist of gender and grades (7 & 8). Five students had an 'unknown' grade assigned and were removed from the computation shown in Table T. School schedule consists of seven class periods, with students generally switching classrooms between periods, in addition to two non-overlapping lunch periods. School hours were 8:25am to 3:15pm on each day.

Data was similar to the SocioPatterns datasets, but collected using Wireless Ranging Enabled Nodes (WRENs). WRENs were distributed to students in their first-period classrooms and collected before the end of last period on each day. Data resolution is approximately 20 seconds (precisely 20,000 binary milliseconds, which equals 19.53125 seconds; there are 210 = 1024 binary milliseconds per second). The cleanup of the original data can be seen in reference [5]. Three networks were made available in the original work [34], one for each day of collection, and a third containing both days. In our analysis we used the combined network containing both days.

Similarly to the other data sets collected in a school environment (see Sections C.1 and C.2, we expect the metalabel classes here to coherently represent the underlying social structure, as can be seen in Fig Z. However, note that the structure captured by the Louvain algorithm contains many more modules than simply the two grades, but the modularity similarity is still high as the underlying social structure is mostly contained within the two grades (see Table T).

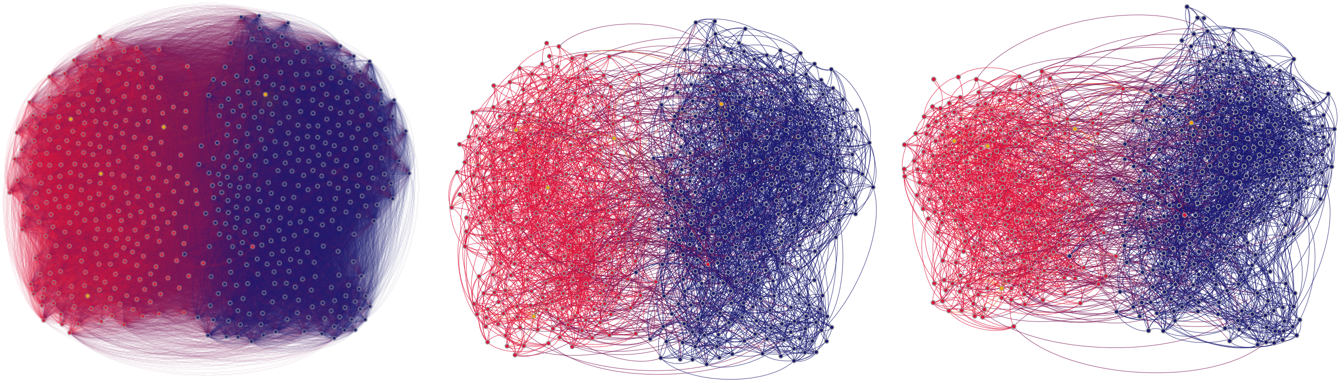

**Fig Z. Contacts in the American Middle School (US-MS).** Distance network, $D(X)$, shown left; backbone, $B(X)$, shown middle and right. Node layout algorithm computed for the original distance network (left and middle) and then recomputed for the backbone (right). Node colors denote student grade: 7$^{th}$ and 8$^{th}$ grades in blue and red, respectively. Students with 'unknown' grade shown in orange.

**Table T. American Middle School (US-MS) measures of module similarity.** Students with 'unknown' grade were removed prior to computation. Values for random subgraphs based on 100 instances.

| | | Original | Metric | Threshold | Random |
|---|---|---|---|---|---|
| | *m* | | | | |
| | Metalabels | 2 | - | - | - |
| | Louvain | 6 | 8 | 11 | 30.49±2.03 |
| | Infomap | 11 | 43 | 51 | 30.49±2.03 |
| Louvain | $y_{AB}$ | | | | |
| | Metalabels | 0.56 | 0.48 | 0.42 | 0.25±0.01 |
| | Original | - | 0.62 | 0.57 | 0.36±0.01 |
| | $J_{A\to B}/J_{B\to A}$ | | | | |
| | Metalabels | 0.61/0.29 | 0.53/0.22 | 0.40/0.16 | 0.11±0.01/0.05±0.01 |
| | Original | - | 0.54/0.45 | 0.49/0.35 | 0.14±0.02/0.08±0.01 |
| | $h_{A\to B}/h_{B\to A}$ | | | | |
| | Metalabels | 0.61/0.31 | 0.67/0.42 | 0.73/0.41 | 0.89±0.02/0.57±0.06 |
| | Original | - | 0.45/0.43 | 0.47/0.38 | 0.77±0.02/0.57±0.03 |
| | *CluSim* | | | | |
| | Metalabels | 0.42 | 0.35 | 0.24 | 0.06±0.01 |
| | Original | - | 0.48 | 0.38 | 0.08±0.01 |
| | *Adjusted Rand Index* | | | | |
| | Metalabels | 0.37 | 0.28 | 0.20 | 0.03±0.01 |
| | Original | - | 0.48 | 0.38 | 0.04±0.01 |
| Infomap | $y_{AB}$ | | | | |
| | Metalabels | 0.56 | 0.48 | 0.42 | 0.25±0.01 |
| | Original | - | 0.62 | 0.57 | 0.36±0.01 |
| | $J_{A\to B}/J_{B\to A}$ | | | | |
| | Metalabels | 0.61/0.29 | 0.53/0.22 | 0.40/0.16 | 0.11±0.01/0.05±0.01 |
| | Original | - | 0.54/0.45 | 0.49/0.35 | 0.14±0.02/0.08±0.01 |
| | $h_{A\to B}/h_{B\to A}$ | | | | |
| | Metalabels | 0.61/0.31 | 0.67/0.42 | 0.73/0.41 | 0.89±0.02/0.57±0.06 |
| | Original | - | 0.45/0.43 | 0.47/0.38 | 0.77±0.02/0.57±0.03 |
| | *CluSim* | | | | |
| | Metalabels | 0.42 | 0.35 | 0.24 | 0.06±0.01 |
| | Original | - | 0.48 | 0.38 | 0.08±0.01 |
| | *Adjusted Rand Index* | | | | |
| | Metalabels | 0.37 | 0.28 | 0.20 | 0.03±0.01 |
| | Original | - | 0.48 | 0.38 | 0.04±0.01 |

**Table U. American Middle School (US-MS) metric backbone statistics.**

| | *social* | *individual time* | *experiment time* |
|---|---|---|---|
| $D(X)$ | | | |
| Nodes | 591 | 591 | 591 |
| Edges | 56,867 | 56,867 | 56,867 |
| $B(X)$ | | | |
| Metric edges ($s_{ij} = 1$) | 3,521 (6.19%) | 3,390 (5.96%) | 3,402 (5.98%) |
| Semi-metric edges ($s_{ij} > 1$) | 53,346 (93.81%) | 53,477 (94.04%) | 53,465 (94.02%) |

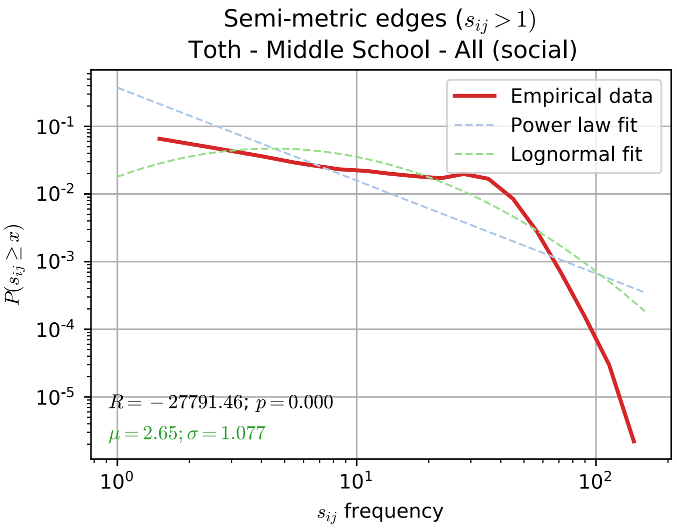


**Fig AA. Distribution of semi-metric distortion values in the American Middle School (US-MS) network.** Log-binned distribution of semi-metric distortion values ($s_{ij} > 1$) for the $\sigma = 93.18\%$ of semi-metric edges in the American Middle School (US-MS) network. Both a log-normal ($\langle s_{ij} \rangle = 2.65$; SD=1.077) and a power law fit are shown; a comparison between the two favours the former as a better representation of the data. Data fitted using the 'powerlaw' python package [3, 4].

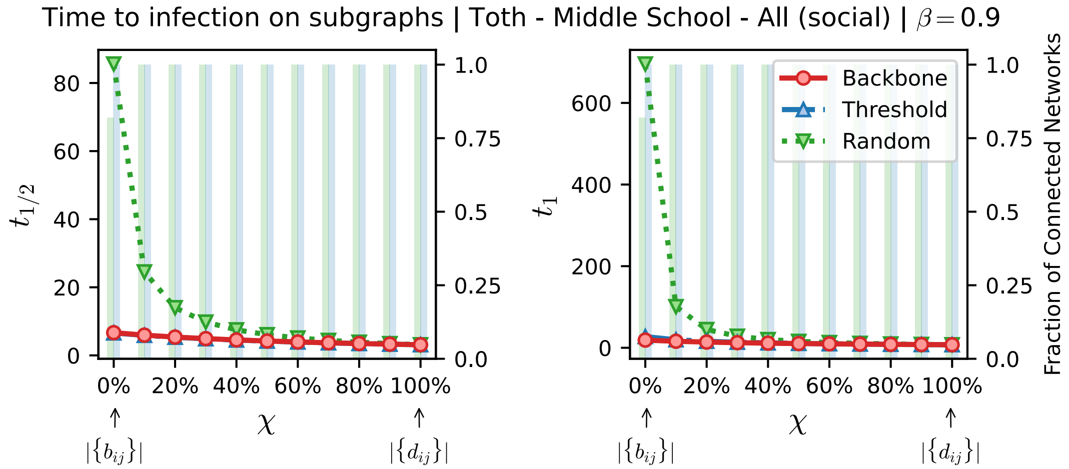


**Fig AB. Time to infection using the metric backbone, threshold, or random subgraphs of the American Middle School (US-MS) network.** The horizontal axis denotes $\chi$, a parameter to sweep the proportion of edges of the original network that are included in the subgraphs analyzed. When $\chi = 0\%$ (leftmost value on axis) we have the metric backbone subgraph, or threshold and random subgraphs with the same number of edges as the metric backbone (i.e. $|\{b_{ij}\}| = \tau(D).|\{d_{ij}\}|$, per eq. 6). As $\chi$ increases, edges from the original network that are not on the backbone or same-size threshold and random subgraphs, are progressively added until the original network itself is reached at $\chi = 100\%$. (Left panels) denotes the time for half of the population to be infected, $t_{1/2}$. (Right panels) denotes the time for all nodes in the network to be infected, $t_1$. The green and blue bars, quantified against the right vertical axis in each panel, denote the fraction of networks in threshold and random baseline ensembles that are connected for a given $\chi$. Non-connected networks are discarded to compute the spreading times. For the simulations shown, the spreading parameter was set as $\beta = 0.9/p_{max}$ where $p_{max}$ is the largest proximity weight of the original network.
